# Towards reliable quantification of cell state velocities

**DOI:** 10.1101/2022.03.17.484754

**Authors:** Valérie Marot-Lassauzaie, Brigitte Joanne Bouman, Fearghal Declan Donaghy, Laleh Haghverdi

## Abstract

A few years ago, it was proposed to use the simultaneous quantification of unspliced and spliced messenger RNA (mRNA) to add a temporal dimension to high-throughput snapshots of single cell RNA sequencing data. This concept can yield additional insight into the transcriptional dynamics of the biological systems under study. However, current methods for inferring cell state velocities from such data (known as RNA velocities) are afflicted by several theoretical and computational problems, hindering realistic and reliable velocity estimation. We discuss these issues and propose new solutions for addressing some of the current challenges in consistency of data processing, velocity inference and visualisation. We translate our computational conclusion in two velocity analysis tools: one detailed method *κ*-velo and one heuristic method eco-velo.

**Author summary:** Single cell transcriptomics has been used to study dynamical biological processes such as cell differentiation or disease progression. An ideal study of these systems would track individual cells in time but this is not directly feasible since cells are destroyed as part of the sequencing protocol. Because of asynchronous progression of cells, single cell snapshot datasets often capture cells at different stages of progression. The challenge is to infer both the overall direction of progression (pseudotime) as well as single cell specific variations in the progression. Computational methods development for inference of the overall direction are well advanced but attempts to address the single cell level variations of the dynamics are newer. La Manno et al. [1] proposed that simultaneous measurement of abundances of new (unspliced) and older (spliced) mRNA in the same single cell adds a temporal dimension to the data which can be used to infer the time derivative of single cells progression through the dynamical process. State-of-the-art methods for inference of cell state velocities from RNA-seq data (also known as RNA velocity) have multiple unaddressed issues. In this manuscript, we discuss these issues and propose new solutions. In previous works, agreement of RNA velocity estimations with pseudotime has been used as validation. We show that this in itself is not proof that the method works reliably and the overall direction of progression has to be distinguished from individual cells’ behaviour. We propose two new methods (one detailed and one cost efficient heuristic) for estimation and visualisation of RNA velocities and show that our methods faithfully capture the single-cell variances and overall trend on simulation. We further apply the methods to a dataset of developing mouse pancreas and show how the method can help us gain biological insight from real data.

## 1 Introduction

Single cell transcriptomics has facilitated the study of asynchronous cellular processes such as cell differentiation in the high-dimensional gene expression space. Development of computational methods for extracting temporal information from snapshots of the system has attracted much attention in recent years. The output of these methods is typically a pseudo-temporal ordering of cells, representing their progression along the (deterministic) path of directed differentiation. However, this ordering does not reflect the intrinsic stochastic characteristics of the process and leaves several biologically interesting questions unanswered. Can cells go back along de-differentiation paths? If yes, how far and how likely is that? How strong is the stochastic component of the dynamics compared to the deterministic directed part? Answering these questions would allow quantification of cell fate plasticity in different transcriptional regions.

RNA velocity, first proposed by [1], was a breakthrough towards obtaining a more complete description of the dynamics of cell differentiation. Simultaneous measurement of abundances of nascent unspliced and mature spliced mRNA in single cells adds a temporal dimension to the collected data which can be used to infer the temporal motion of single cells in transcriptomic space. A later method, scVelo [2], further advanced the concept by solving the transcriptional dynamics of splicing kinetics and velocity inference. Other extensions included additional temporal layers of gene regulation such as protein levels [3] or chromatin accessibility [4] to the unspliced and spliced mRNA levels to extract further information on cell state dynamics. Recently, there have also been advancements in using cell state velocities to study the degree of cell plasticity [5]. For all these methods, it is important to first ensure robust and reliable estimation of single cell velocities. Ideally, the estimated velocities should capture both the overall course in the population as well as the single-cell specific (stochastic) part of the dynamics. However, reliable inference of cell state velocities is still impeded by multiple computational issues. Some weaknesses in current velocity visualisation approaches, as well as challenges in inclusion of genes with multiple dynamics, have been pointed out in [1, 2] and [6]. Another issue on scale invariance of gene-wise velocity components was described in more detail in [7]. Current methods either do not address this scale invariance issue or address it incompletely through computationally costly workarounds. Moreover, there are several inconsistencies in the current methods’ processing pipeline and the stochastic part of the dynamics is lost through multiple layers of data imputation and smoothing. In parallel to this study, [8] also investigates some of the current problems in existing methods for RNA velocity estimation.

In the current manuscript, we revise several steps of velocity estimation and visualisation. We also adjust the processing pipeline to be consistent with downstream velocity estimation. We propose two different approaches for estimation and visualisation of RNA velocities. In *κ*-velo we solve for the genewise rate parameters as done in scVelo but propose a new approach which allows us to estimate the relative magnitude of velocity components across genes, hence solving the scale invariance issue. We also present a novel visualisation method for the inferred velocities. In addition, we propose a second approach, eco-velo, as a heuristics method which bypasses several cumbersome, computationally costly and stochasticity killing steps used by other available methods.

## 2 Methods

### 2.1 Dynamical inference

Building high-dimensional cell state velocities as vector sums of their gene-wise components (as is the current practice) requires careful handling of two major issues: ambiguity of the time scales and the relative scaling between different velocity components. In this section, we discuss current problems in state-of-the-art velocity estimation approaches and introduce our novel *κ*-velo and eco-velo approaches.

#### 2.1.1 Time scale over which average cell state velocities are reported

In the physical world, we can only measure average velocities in a given time interval Δ*t*. As Δ*t →* 0 measured velocities get closer to instantaneous velocities, which are impossible to measure directly. When adding multiple velocity components one would ideally need to measure all gene-wise displacement components 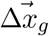 in the same interval Δ*t*. Mathematically:

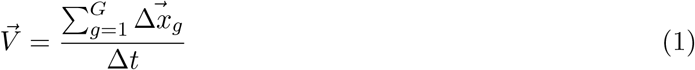

where 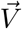 is the G-dimensional velocity vector and Δ*t* is the same for all genes. However, in the RNA velocity framework (even without scale invariance problem discussed in the next subsection) we use:

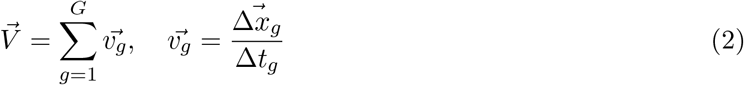

the result of which is different from Equation 1 for non-smooth expression dynamics. Using a different Δ*t*_*g*_ for each gene *g*, raises an immediate question: to which time interval does the average cell state velocity 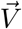 calculated from Equation (2) correspond? Obscurity in the physical meaning of velocities calculated as such is more pronounced when including genes with noisy expression dynamics, e.g. bursting genes. For noisy genes, velocities reported over different time scales will change (Fig. S1). For such genes, it would be interesting to experimentally measure velocities at multiple time scales. This could help us better understand the extent of cell fate plasticity. One would expect to see more variance in the direction of individual cell velocities reported in small time scales, whereas velocities over sufficiently large time scales would better align with the pseudotemporal direction of differentiation.

#### 2.1.2 Scale invariance of gene-wise velocity components

According to the RNA velocity formalism:

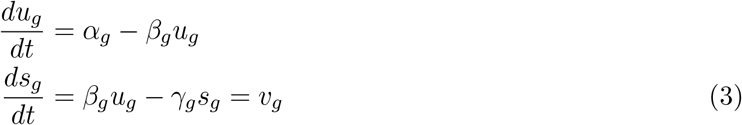

where *u*_*g*_ and *s*_*g*_ represents the number of unspliced and spliced counts for gene *g. α*_*g*_, *β*_*g*_, *γ*_*g*_ represent transcription rate, splicing rate and mature mRNA degradation rate respectively. *v*_*g*_ represent the instantaneous velocity component of gene *g*. It has been shown that there is a scaling invariance in the solution for the parameters that one can infer from the unspliced-spliced (u-s) phase portrait, meaning that if (*α*_*g*_, *β*_*g*_, *γ*_*g*_) is a solution, (*κα*_*g*_, *κβ*_*g*_, *κγ*_*g*_) is also a solution for any *κ*. Thus to get a valid high-dimensional 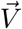, the gene-specific solutions to Equation 3 do not suffice and we need to know the real scaling factor *κ*_*g*_ for each gene:

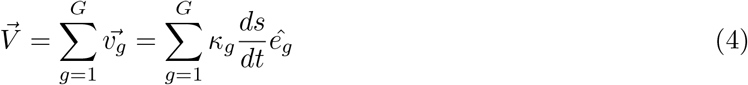

where 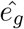 represents the unit vector for gene g.

To overcome the scale invariance, velocyto assumes *κβ* = 1 (i.e. same splicing rate) for all genes. scVelo, however, assumes a constant prior time (equal to 20 hours) for a full cycle (i.e. turning on, reaching stationary state and turning off) on the u-s phase portrait for all genes in order to estimate gene-wise latent time for each cell. It then assigns a “gene-shared latent time” to each cell using a multi-step ad-hoc voting method that combines the fitted latent times from multiple high-likelihood genes. To circumvent the scale invariance in *κ*-velo, we suggest a more straightforward approach, which is to use a proxy for gene-wise typical travel time from the expression level in cell *i* to the expression level of cell *j*. In eco-velo we take a different approach, which, similar to velocyto, relies on strongly simplifying assumptions on the kinetic rate parameters.

#### 2.1.3 First approach: *κ*-velo

Our first approach addresses the scale invariance problem by using a proxy of time between two measurements which allows for estimation of the scaling factors of genes. In this subsection, we drop the gene-wise indices *g* and use cell-wise indices as we address the scaling factor *κ* for one gene at a time.

Let Δ*t*_*ij*_ be a measure of time that can be used to relate time between two states *i, j* across genes, with *i* before *j* in time. Consider one gene with true parameters of reaction rate *θ* = (*κα, κβ, κγ*) and recovered parameters *θ* = (*α, β, γ*) and *u*_*i*_, *u*_*j*_ the measured unspliced counts for cells *i, j*. Note that for that gene, *i, j* need to be in the same state of transcriptional induction or repression because the speed of genes is only measurable during transcriptional change, i.e. outside of steady-state. If the cells spend time in steady state, the change in transcriptional state will not be proportional to the distance in time, which is why we only consider cells in the same state.

The solution of Equation 3 for u counts is 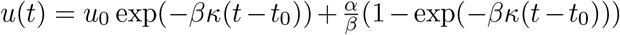, where *u*_0_ is the initial condition of the u counts at time *t*_0_. This yields for two measurements from cell *i* and *j*:

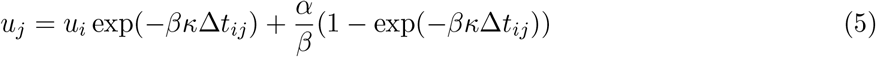

Solving for *κ*Δ*t*_*ij*_ we get:

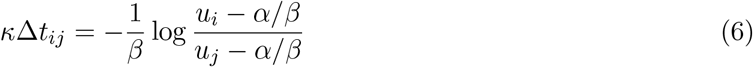

One can solve the equation of *s*(*t*) similarly but we focus on *u*(*t*) as it depends on one less parameter (see Supplementary Note S1).

Since we do not have a true measure of Δ*t*, we propose that cell densities, i.e. number of cells that are captured between two cell states, can be used instead. In snapshot data, the probability of capturing cells in a specific region of the expression space is proportional to the time cells typically spend in that region. Therefore cell density is inversely proportional to the average velocity in any expression region. This relation may approximate several single-cell datasets, although it is disturbed in presence of non-uniform cell proliferation and death rates as well as biased sampling of cell types (e.g. enrichment of specific cell types) in the experimental set up. In those cases, it is also possible to design other proxies for estimation of Δ*t*_*ij*_, such as the diffusion distance.

To recover *κ* from Equation 6, we randomly sample pairs of cells *i, j* that are in the same transcriptional state. We compute *d*(*i, j*), the number of cells between the states, and *f* (*i, j*), the right part of Equation 6 for each pair. For pairs in transient state, *f* (*i, j*) should grow linearly with *d*(*i, j*), while if either (or both) cell is in steady-state, *f* will be smaller than expected from Equation 6. Plotting *d* on the *x*-axis and *f* on the *y*-axis, will produce a parallelogram of which the left slope equals *κ*. To recover *κ*, we fit a parallelogram to the points such that the area is minimal while maximising the number of points in the parallelogram (Note S2, Fig. S2).

We note that our approach addresses the scale invariance problem but, like other existing methods, it does not resolve the ambiguity in the time scale over which cell state velocities are reported as we use different Δ*t*_*ij*_s (different pairs of cells) for inferring a common *κ* for all of those times. Therefore exclusion of noisy genes is important. Noisy genes also cause issues during the assignment of cells to the state of transcriptional induction or repression and thus impair the estimation of kappa. For these two reasons, we exclude noisy genes.

After determination of the gene-wise *κ*_*g*_, we call the high-dimensional, correctly scaled parameters Θ = (*A, B*, Γ). For calculating the velocity for cell *i*, we use 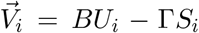, where *U*_*i*_ and *S*_*I*_ respectively represent the G-dimensional u-s counts in cell *i*. Note that in the scVelo approach, fitted latent time assignment *t*_*i*_ for cell *i* is used to impute its u-s coordinates *u*(*t*_*i*_) and *s*(*t*_*i*_), which is then used as 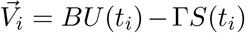. Using imputed u-s coordinates, as scVelo does, introduces an additional layer of data smoothing that removes biologically relevant variance.

#### 2.1.4 Second approach: eco-velo

Starting from Equation 3, for the change of the spliced counts of gene g over Δ*t* we can write:

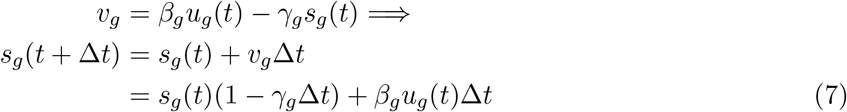

By fixing Δ*t*_*g*_ = 1*/γ*_*g*_ we get:

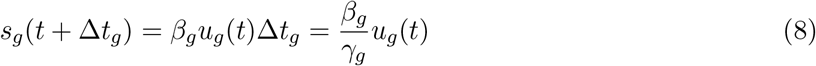

This means that knowing *β*_*g*_*/γ*_*g*_ is sufficient to estimate the cell state displacements over Δ*t*_*g*_. If we further assume all genes have a similar *β* and *γ*, we can conclude spliced counts at time *t* + 1*/γ* will look like the u counts of an earlier time point *t* as *s*_*g*_(*t* + Δ*t*_*g*_) = *u*_*g*_(*t*).

This leads to another level of simplification that turns out very handy as a heuristic velocity estimation from u and s counts, where we can find cell state displacements by mapping U to S. We do so by searching for the nearest neighbor (NN) of U in S that is also within the first *k* nearest neighbors of S in U. We call these pairs mutual nearest neighbors (MNNs). Note that not every point needs to have a MNN. The velocity arrow then goes from a cell’s position in S space to the first MNN of that same cell’s U space in S. Here, u and s counts can be used directly for estimating cell state velocity directions without any need for strict gene set selections, smoothing and parameter fitting, all of which takes a lot of computational power and kills the interesting (biologically meaningful) variance.

Similarly to velocyto and scVelo, this approach does not account for the time scale (here 1*/γ*_*g*_) of the observed displacements being different for different genes. However, the assumption of similar *β* and *γ* for all genes has been shown to hold for some systems. In Fig. 1e from the original RNA velocity paper [1], we see bulk RNA-seq measurements in the mouse liver over a time course of the circadian cycle where unspliced mRNAs are predictive of spliced mRNA at the next time point.

**Figure 1:**
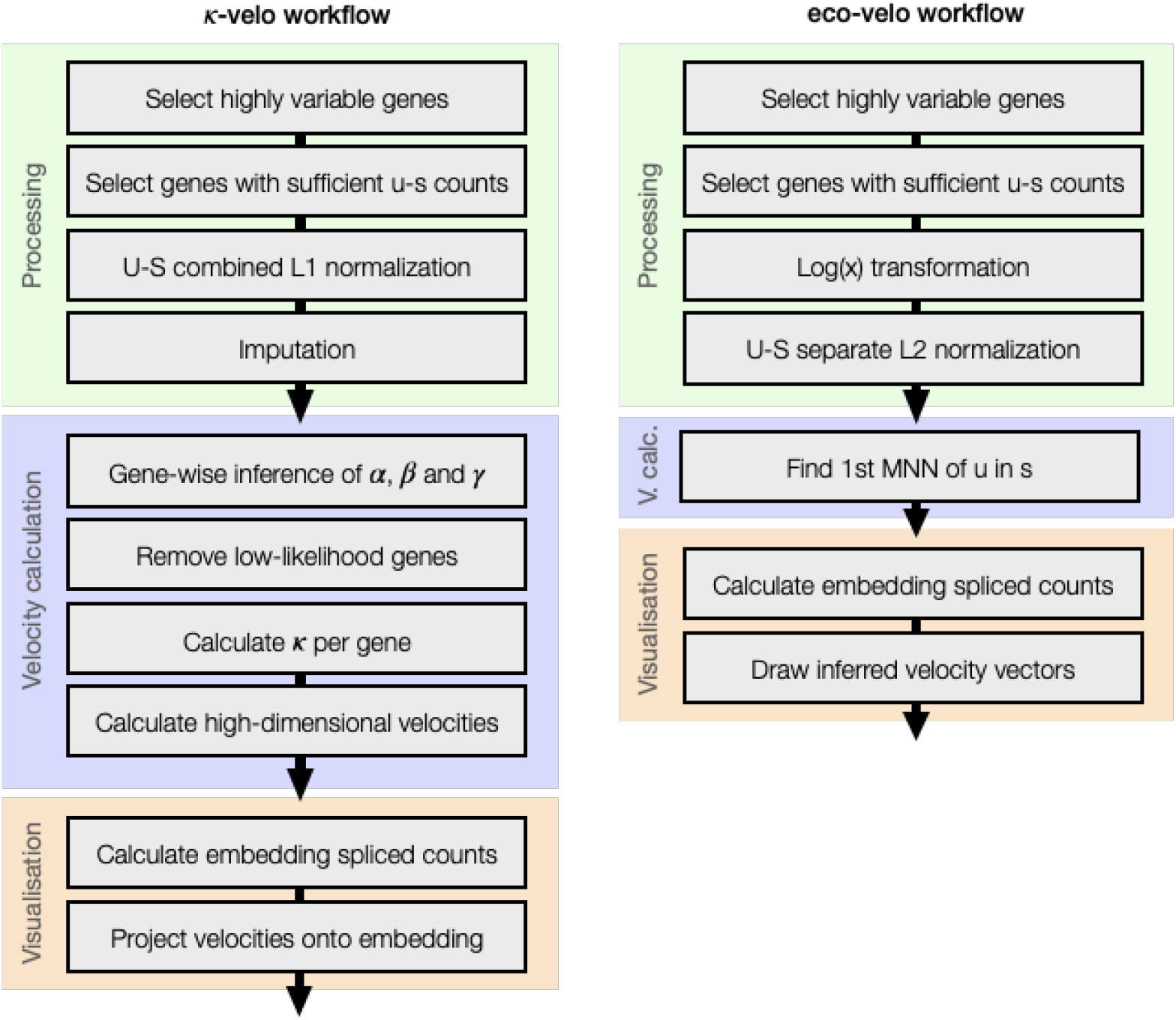
An overview of RNA velocity analysis steps in the *κ*-velo and eco-velo workflow.

### 2.2 Visualisation

La Manno et al. [1] suggested using projection of the end of the velocity vectors (*s* + *v*Δ*t*) on an embedding of the spliced counts (*s*). While projection using principal component analysis (PCA) (Note S3) is the most accurate low-dimensional representation of cell state velocities, it usually does not capture the full complexity of differentiation manifolds with several subpopulations in high-dimensional gene space. Projection of the velocities onto non-parametric nonlinear embeddings (which do not have gene-defined axes) is more challenging. To work around this difficulty, velocyto projects the velocities in a direction relative to the neighbouring cells. This is done by computing a transition probability matrix *P* containing probabilities of cell-to-cell transitions in accordance with the velocity vector: 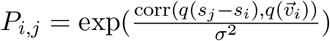 with *σ* the kernel width parameter, 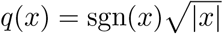 a variance-stabilising transformation and corr() the Pearson correlation coefficient. The matrix is row-normalised so that ∑_*j*_ *P*_*ij*_ = 1. Given *n* observations and *y*_*i*_ a vector of the positions of cell *i* on an embedding, the projected velocities are estimated as:

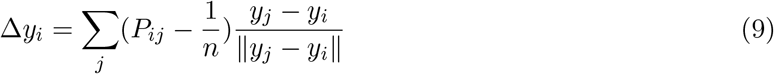

To project the velocities, scVelo uses a similar approach to velocyto but with a slightly different *P* matrix based on cosine similarity cos ∠() in 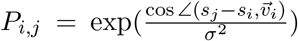. A vector summation as in Equation 9 used in velocyto and scVelo is questionable as the direction of several neighbouring cells can be correlated on the low dimensional embedding. Therefore 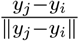 does not in general provide an orthonormal basis. As a result, this approach changes the direction of the velocity vectors as well as their magnitude. Another issue with this approach is that it removes a considerable part of the variation among individual cell velocities by projecting them on the direction of nearest neighbours only. Lastly, the size and shape (i.e. extension along the differentiation path versus its width) of the considered neighbourhoods will affect the velocity directions. For example if different subpopulations share expression similarity with the direction of the velocities, these multiple targets can rotate the direction of projected velocities. Another example is if the neighbourhood size is larger than the distance to the ultimate trajectory target. Here too the direction of projection can rotate away from the intended target.

We argue that projection by PCA is the only projection method where the arrows are a true representation of the high-dimensional velocity vector. To deal with more complex data manifolds, we propose a new approach for nonlinear projection that is more faithful to the true cell state velocities than the current nonlinear embedding practices. For *κ*-velo, we use linear and nonlinear projection methods. For eco-velo, visualisation of velocities is integrated within the inference of the velocities method and hence does not require visualisation by projection.

#### 2.2.1 Projection onto nonlinear embedding (visualisation for *κ*-velo)

We project the points of the velocity arrows (as test set data points) onto an existing nonlinear embedding, such as diffusion maps [9], t-distributed stochastic neighbor embedding (t-SNE) [10] and uniform manifold approximation and projection (UMAP) [11], of the initial cell positions (training set). To do this, we first compute a transition probability matrix *P*_*n*1*×n*1_ between all pairs of cells *i, j* in the training dataset containing *n*1 cells. We then solve for a matrix *W* : ℝ^*n*1^ *→* ℝ^*k*^ that transforms the *P* matrix into the pre-trained *k*-dimensional embedding, i.e. *PW* = *Y*_*n*1*×k*_. A new 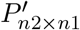 is then calculated between the new (test) data points and the training set. The projection of the new points is given by *P* ^*t*^*W*. (Note S4).

Such a nonlinear projection approach is also known as Nyström projection and has already been used for single cell data integration applications [12, 13]. In fact, what we propose here is an adaptation of the Nyström method [14] for visualisation of the velocity vectors (see also Note S4).

It is important to note that unlike parametric PCA projections, nonlinear projection approaches as described above are only valid if the test data is close enough to the data points in the training set. Hence, extrapolation for test data to expression regions which have not been sampled in the training set is not possible. This is even more crucial for t-SNE and UMAP (versus diffusion maps) which may not preserve the global continuity of the intrinsic data manifold. Closeness of the point of the velocity arrows to the data manifold of the base of the arrows can be guaranteed when velocities are small enough.

The embedding of current cell states onto which the future states are projected is calculated on the same gene space as the space used for velocity recovery. This ensures that the embedding only represents a space that can be spanned by following the calculated velocity directions. We further calculate the embedding on the same space used for parameter recovery. This means that if we use imputed counts for parameter recovery, we calculate the low-dimensional embedding on those imputed counts.

#### 2.2.2 Visualisation for eco-velo

We identify the first mutual nearest neighbour (MNN) [15] of U and S for every cell, which we use to visualise the velocities on a low-dimensional embedding of the spliced counts. We simply draw an arrow starting from the position of a cell on the embedding to the coordinates of its first MNN on the same embedding. That means that our velocity arrows point from *s*_*i*_ to *s*_*j*_, where *s*_*j*_ is the first MNN of *u*_*i*_ for cell *i*. These arrows corresponding to a relatively large Δ*t* in which all current spliced counts in the cell would be degraded. For ease of visualisation and obtaining an un-cramped map without intersecting cell velocities, we then scale all velocities by the same factor so that the arrows only point in the direction of the point and not all the way to the future state.

### 2.3 Processing

Before calculating the velocities, single-cell RNAseq datasets are preprocessed (aligning the reads and counting numbers of unspliced and spliced reads) and processed (filtering, normalization, etc.). Both the *κ*-velo and the eco-velo workflows start with processing raw U and S count matrices. Since the methods are based on different assumptions, the processing steps differ per method. Below, we will describe the processing protocol for both approaches.

#### 2.3.1 Processing pipeline of *κ*-velo

To reduce the number of dimensions of the dataset, we select only genes with high variability. Variability is calculated on the spliced counts using analytic Pearson residuals [16]. We then filter genes with extremely low u or s counts because we want to focus only on genes with significant velocity signal. After gene filtering, the counts in each cell are size-normalised. The size of a cell is represented by its u and s counts together. Therefore, the counts in each cell are normalised using a size factor that is calculated on the total counts of u and s. To recover the dynamics, the noise in the u and s counts has to be reduced. As such, all counts are imputed by averaging the counts of each cell’s nearest neighbours. The nearest neighbours for each cell are found in PCA space calculated on scaled s counts. For a more detailed description of each step see Note S5 and Fig. S3.

#### 2.3.2 Processing pipeline of eco-velo

Similar to *κ*-velo processing, the eco-velo workflow starts by filtering the dataset for genes with high variability and sufficient u and s counts. After this, all non-zero counts are log-transformed and both count matrices are normalised separately. Here, we deviate from the *κ*-velo protocol, because u and s counts are treated as separate entities. Following standard MNN protocols [15], the counts are L2 normalised.

### 2.4 Overview of the workflow for *κ*-velo and eco-velo

Both the *κ*-velo and the eco-velo workflows consist of three main steps: processing, velocity calculation and visualisation (Fig.1). First the data is processed as described in Section 2.3. In *κ*-velo, after processing, scVelo is used to recover the parameters *α, β* and *γ* of all genes in the dataset. For downstream velocity analysis, only genes with a likelihood above a certain threshold are used. All other genes are filtered out to reduce the technical noise caused by poorly recovered or noisy genes. We then recover the scaling factor *κ* for each gene, which is used to scale the parameters. Using the scaled parameters, a high-dimensional velocity vector is calculated for each cell. To visualise the cells and velocities, we compute an embedding (e.g. PCA, UMAP) using the processed (i.e. filtered, normalised and imputed) and scaled s counts. Lastly, the velocities are projected onto the embedding.

The eco-velo workflow includes fewer steps. After processing, the u counts are used to find the mutual nearest neighbour of each cell in S space. The embedding is calculated using processed (i.e. filtered and normalised) s counts and velocities are projected onto the embedding by connecting each cell to its first nearest neighbour.

### 2.5 Simulation data

For the simulation, we randomly sampled *g* log-normally distributed parameters of reaction rates, scaled by a scaling factor *κ*: *θ* = (*κα, κβ, κγ*). The true time of the *n* observations is sampled from a uniform distribution. The u and s counts are simulated following *u*(*t*), *s*(*t*) (Note S6) with added random normal noise. We simulate the data such that the time of activation of each gene’s transcription is inversely proportional to the gene’s speed. This means that the fastest genes are only active towards the end of the differentiation trajectory. The resulting differentiation trajectory has high plasticity at the beginning when most genes are not yet commited to change and more deterministic dynamics at the end of the trajectory. See Note S6 for a more detailed description.

### 2.6 Real data

We demonstrate the performance of *κ*-velo and eco-velo on two different datasets and compare them with the state-of-the-art scVelo. The first dataset is a subset of the pancreatic endocrinogenesis dataset [17] and the second is a subset of the murine gastrulation dataset [18]. In this manuscript, our analysis starts from the U and S count matrices, which were originally analysed in [2, 6].

For *κ*-velo, we selected the top 5000 highly variable genes (HVGs). From those, we selected only genes with a maximum u and s count above 3, giving us 716 genes. Both the U and S matrix were L1 normalised, using the sum of the u and s counts as size vector. NNs for each cell were found using the first 15 dimensions of a recalculated PCA embedding. The 30 NNs were used to average both the u and s counts in each cell (imputation). After recovery of the dynamics, only the genes with a likelihood above 0.4 (117 genes) were kept for velocity analysis.

For eco-velo, we selected 2000 HVGs. 395 of the HVGs had a maximum u and s count above 3. The U and S matrices were L2 normalised using separate size vectors. For the selection of MNNs, we set the neighbourhood size *k* = 50.

The erythroid lineage of the murine gastrulation was processed using two different pipelines. First, we used the exact processing pipeline reported in [6]. Then we used our processing pipeline where we selected the top 5000 HVGs. We selected genes with a maximum u and s count above 3 and applied u-s combined L1 normalisation. Lastly, we imputed the counts (PCA embedding with 15 dimensions; 30 NNs).

## 3 Results

In this section, we first demonstrate the artefacts of scVelo’s velocity projection on simulation data with known cell state velocities (i.e. no velocity inference step involved) and compare scVelo to visualisation with linear and nonlinear projection methods. We then compare our *κ*-scaled velocities with the velocities returned by scVelo on simulation. Afterwards, we compare the results of our complete (i.e. velocities inference and visualisation) *κ*-velo and eco-velo workflow with those from scVelo on real data. In the last section, we show computational experiments on real data which support the design of the processing steps we propose and use in this manuscript.

### 3.1 Visualisation by linear and nonlinear projection methods faithfully represents true velocity directions

Ideally, a visualisation of cell state velocities should be faithful to the projection of the high-dimensional direction of velocity vectors as well as their magnitude (speed of change) onto the existing data embedding. It should also preserve local variations, representing fluctuations of the dynamics and cell plasticities. To assess these points, we compare existing RNA velocity visualisation methods with ours, on simulated data where the true high-dimensional velocities are known and do not need to be inferred. We simulate a single cell dataset with cells following a hidden true time with high plasticity at the beginning and faster commitment to fate towards the end of the trajectory (Section 2.5). Projection of the velocities on a PCA embedding (Fig. 2A) reliably represents all these aspects. scVelo’s velocity projection on the same PCA embedding (Fig. 2B) smooths over the biologically interesting variation and removes the information of speed of change. Our nonlinear projection method (Fig. 2C) captures the expected cell to cell variation, as well as the direction and speed of the simulated velocities on a t-SNE embedding of the data (Fig. 2D and S4A-B). Our projection method’s results for diffusion maps and UMAP are included in Fig. S4.

**Figure 2:**
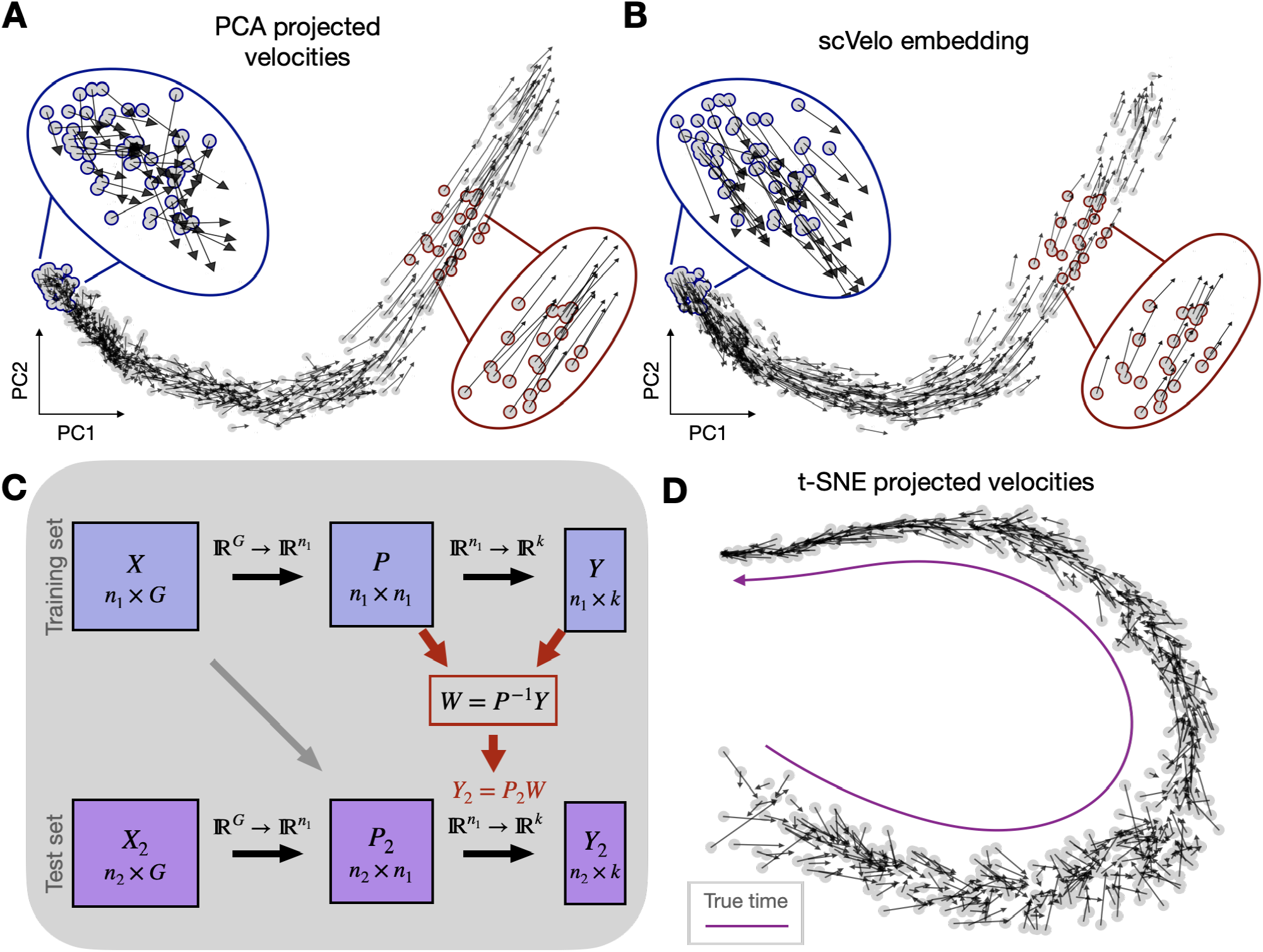
Visualisation of simulated velocities with linear and nonlinear projection methods. **A**. Velocities projected on PCA embedding. The blue outline puts forward a region of high plasticity and the red outline shows a low-plasticity, high-velocity region. The arrows in the PCA embedding capture both the plasticity and length of velocities. **B**. Velocities projected on PCA embedding by scVelo. scVelo smoothes over variance, thereby losing the information on cell plasticity as illustrated in the cells outlined in blue. scVelo also loses the information of vector length as shown in the cells outlined in red. **C**. We propose a method to project the future states (test set) onto an existing nonlinear embedding of the current states (training set). We recover *W*, a transformation of the transition probability matrix *P* into projected cell states *Y*. We then apply the same transformation on *P*_2_, the transition probability matrix from training set to test set. This gives us the projected future states *Y*_2_. **D**. Velocities projected by projection method (c) shown on t-SNE.

### 3.2 *κ*-velo recovers simulated velocities

To ensure that the high-dimensional velocity vector points in the right direction we need to address the scale invariance of gene-wise velocity components (as discussed in Section 2.1). We develop *κ*-velo, a method that uses the cell densities as a proxy of time spent in a specific region of the expression space (Fig. 3B). This time proxy allows us to relate velocities across genes. To validate our method we simulate reaction kinetics following randomly sampled parameters scaled by a factor *κ* varied between 1 and 15. The method recovers the scaling factors (Fig. 3C). Note that the recovery becomes more difficult for higher *κ*. Very fast genes have few or no cells in transient state so in those cases we would need to sample more cells to reliably recover *κ*. The scale of recovered *κ* and true *κ* is still off by a constant factor related to the chosen Δ*t*. However, if all components are scaled by the same factor, the direction of the high dimensional vector is still correct. Because of this, the high dimensional *κ*-velo velocity vector is much closer to truth (Fig. 3D). Consequently, the velocities projected on a PCA embedding are also much closer to truth, both for direction and length (Fig. 3E).

**Figure 3:**
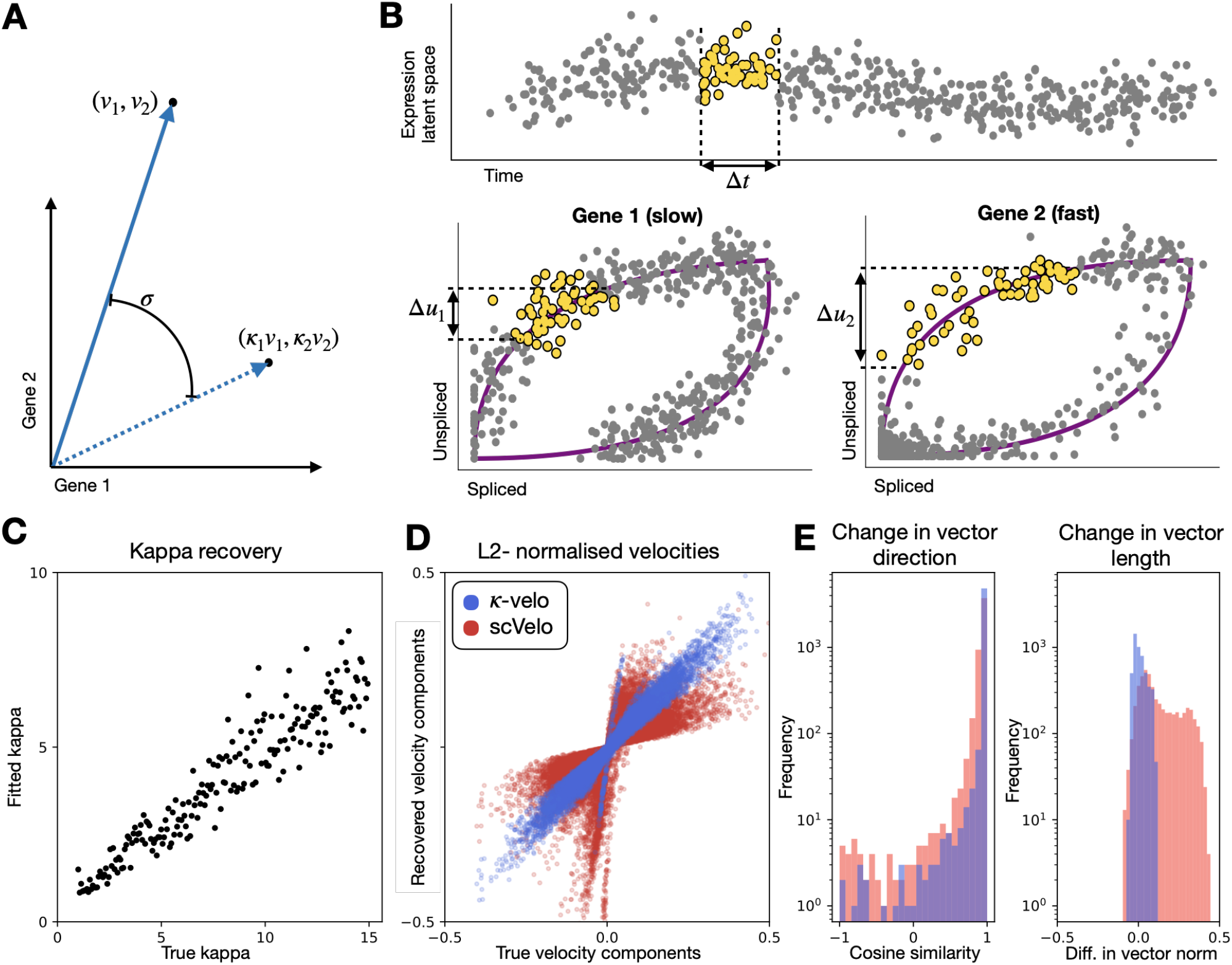
Scaling of gene-wise velocity components. **A**. If the gene-wise velocities are incorrectly scaled the high-dimensional velocity vector will change direction (displacement *σ*). **B**. We propose to use cell densities as a proxy of time. For a same time interval, the displacement in u will be proportional to a gene’s speed. This allows us to relate velocities across genes and solve the scale invariance problem. **C**. To validate *κ*-velo, we simulate splicing kinetics scaled by a scaling factor *κ* and evaluate how well the factors are recovered. **D**. We compare the high-dimensional *κ*-velo and scVelo velocities to the true velocities. One point represents a velocity for one cell for one gene. The high-dimensional velocity vector is L2 normalised for ease of comparison. **E**. The high-dimensional vector is projected on the first two principal components to evaluate differences between true velocities and recovered velocities. We return the change in direction (cosine similarity) and length (difference in vector norm) (Note S7) for *κ*-velo and scVelo. To make the length comparable, here too the vectors are L2 normalised. Note the log-scale for frequency.

### 3.3 *κ*-velo explains cell state plasticities and speed of transcriptional change in pancreas endocrinogenesis

To test whether *κ*-velo’s velocity estimations better capture the different time scales of genes, we apply our method to a dataset of developing mouse pancreas cells sampled at embryonic day 15.5 [17]. The endocrine progenitor cells differentiate into four main fates: alpha, beta, delta and epsilon cells. In previous work, scVelo delineated cycling progenitors and the endocrine cell differentiation. After processing, we get the reaction rate parameters fitted by scVelo and recover the scaled parameters for *κ*-velo. The range of splicing rates returned by scVelo is more than 10000-fold (Fig. 4A and S5). True splicing rates are difficult to determine and different ranges have been reported [19] but none come close to the range reported by scVelo. We report a range of splicing rates close to 30-fold (Fig. 4A), which is closer to expected in the biological context. After scaling, we can distinguish fast and slow genes based on their *κβ*. We find a substantial amount of genes associated with the cell cycle within the fast genes, such as *Nusap1, Adamts1* and *Anxa2*, while slow genes are constantly up- or down-regulated during the whole differentiation trajectory (Fig. 4B). This is consistent with biology as the cell cycle in developing mouse pancreas takes less than a day [20], while pancreatic endocrine cell differentiation starts at embryonic day 9 and goes until 15.5 in the analysed sample. We also find fast genes that are up-regulated during commitment to a cell fate at the end of the differentiation trajectory, such as *Gcg* and *Nnat*.

**Figure 4:**
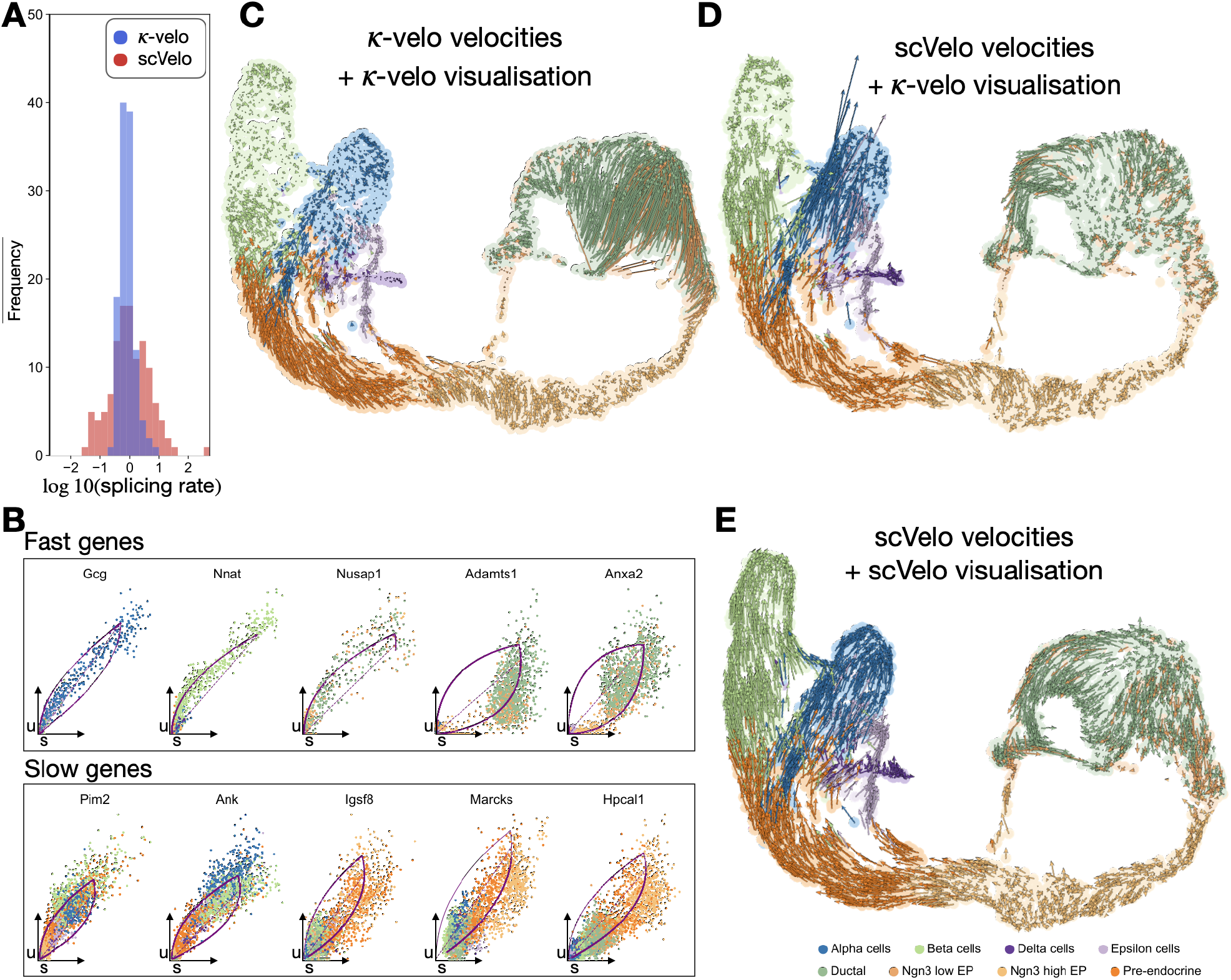
*κ*-velo on pancreas endocrinogenesis. **A**. Range of splicing rate *β* estimated by scVelo (in red) and *κ*-velo (in blue). **B**. Examples of fast and slow genes, selected according to *κβ*. Learned kinetics are shown by purple curves. **C**. Velocities from *κ*-velo projected onto a UMAP embedding using *κ*-velo projection. **D**. Velocities from scVelo projected onto the same UMAP embedding using *κ*-velo projection. **E**. Embedded velocities as returned by scVelo. For ease of comparison, plotting style was matched to (c) and (d).

We display the high dimensional vector field in a UMAP embedding of the data and compare the *κ*-velo velocities (Fig. 4C) to scVelo velocities (Fig. 4D), both projected with *κ*-velo (Fig. S6 shows smoothed velocities on the embedding). The *κ*-velo velocities better capture the differences in speed along the trajectory. Towards the end of the alpha cells, scVelo returns conflicting velocities, which can in parts be traced back to the glucagon-encoding gene *Gcg* that is not fitted according to what would be expected from biology. The scaled velocities better capture the terminal states towards the ends of the fates but there is still some ambiguity. We note that a lot of genes do not display evident kinetics on the u-s phase trajectory, which can cause issues during fitting of reaction rate parameter. One example is given by *Gcg*, where the shape of the trajectory cannot be clearly identified as transcriptional induction or repression. scVelo’s embedding (Fig. 4E) smooths over the velocities, returning a view that appears more consistent with the expected direction of differentiation but not with the actual noisy velocity vectors.

### 3.4 Eco-velo approximates velocity directions using minimal data processing and computation

As a heuristic method that does not require cumbersome recovery of the rate parameters, we introduce eco-velo. By simply taking the unspliced counts as a proxy of a cell’s future state (Fig. 5A), we can skip strict gene set selection, imputation and parameter fitting, all of which are computationally expensive and can kill some of the signal. We validate the model on a simulated dataset (Fig. 5B and S7), where the model recovers the expected flow. We then test eco-velo on the pancreas endocrinogenesis dataset (Fig. 5C and S8 for smoothed velocities). The model clearly delineates the directional flow from progenitor cells to alpha and beta cell fates. eco-velo also captures the high cell plasticities towards the end of the differentiation trajectory seen in Fig. 4C. The final state of epsilon cells is also captured (Fig. S8 smoothed) but the dynamics within the delta cells cannot be resolved. For delta and epsilon cells the issues could arise from trying to capture future states within sparse populations that are transcriptionally close to the more abundant population of alpha cells. Given the strong theoretical assumptions of the model, eco-velo still captures the complex lineages of endocrinogenesis remarkably well.

**Figure 5:**
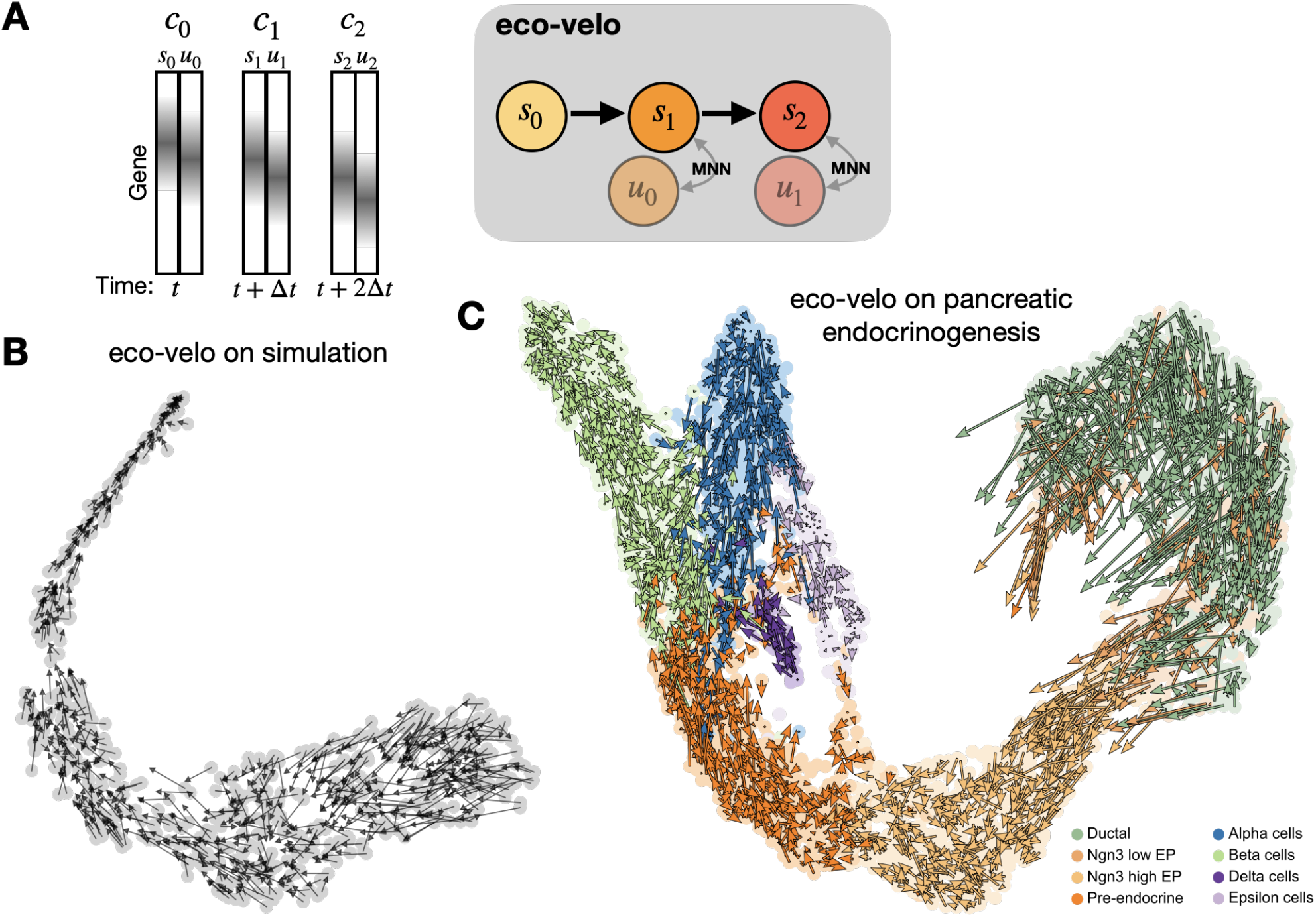
eco-velo as an alternative to computationally costly reaction rate parameter recovery. **A**. Under certain conditions, a cell’s unspliced state will represent the cell’s future spliced state. To infer velocities, we look for the first MNN between a cell’s unspliced counts and other cells’ spliced counts. We draw an arrow from the cell to the identified MNN. **B**. We validate eco-velo on simulation and visualise the resulting velocities on t-SNE. **C**. eco-velo on pancreas endocrinogenesis.

### 3.5 Careful processing prevents introduction of artefacts

To illustrate the importance of processing, we apply our processing pipeline to a dataset of erythroid development during murine gastrulation. Previously, it has been shown that scVelo falsely predicts de-differentiation at the end of erythroid development. This has been attributed to the contribution of genes with multiple rate kinetics (MURK genes) to the velocity calculation [6]. In our processing pipeline, we not only filter for low variability genes but also remove genes with insufficient u and s counts. This step is important as genes with only few u or s counts usually produce unreliable results after imputation (Fig. S9). In the regular scVelo processing pipeline, the count matrices are normalised separately. This separate normalisation introduces artefacts in the u-s portrait (Fig. S10A-B), which can be traced back to variation in the ratio between total unspliced and total spliced counts between per cell types. We found that some of the patterns identifying MURK genes were artefacts of this normalisation (Fig. S10A-B). After normalisation, the counts are imputed by averaging spliced or unspliced counts across neighbouring cells, thereby smoothing the data. Many MURK genes in the original publication were imputed from very low counts and are filtered in our pipeline. Comparing the original processing pipeline to our processing steps, we reduce the number of MURK genes from 98 to 18 (Fig. S10C), correcting most of the false de-differentation.

After recovery of the parameters, our *κ*-velo workflow includes an additional filtering step where we remove genes where the learned parameters do not describe the observed cell states very well. This prevents us from including the (usually noisy) genes for which the recovered parameters are less meaningful (Fig. S11). The calculated velocities for those genes would therefore not accurately reflect true dynamics. We also calculate the low-dimensional embedding on this reduced gene set, so that the embedding only represents space that can be reached by velocities. Interestingly, our complete workflow resolves the de-differentiation in epsilon cells reported in the scVelo paper (Fig. 4E). The results of processing on both datasets emphasise the necessity of careful processing.

### 3.6 Computational efficiency of the methods

We report the runtime on an Intel Core i5 CPU with 2GHz, 4 Cores and 16 GB of RAM. On the pancreatic endocrinogensis dataset with 3696 cells the *κ*-velo workflow takes 4 minutes while the eco-velo worflow takes about 40 seconds.

Full scVelo pipeline on the same dataset takes about 8 minutes.

### 3.7 Data and software availability

All analysed datasets are publicly available. The pancreatic endocrinogenesis dataset is available from the Gene Expression Omnibus under accession GSE132188 [17]. The murine gastrulation dataset is available on the Arrayexpress database (http://www.ebi.ac.uk/arrayexpress) under accession number E-MTAB-6967 [18]. For both datasets the count matrices can be downloaded directly from the scVelo Python implementation (https://scvelo.org) v0.2.4.

All Python scripts and notebooks necessary to reproduce the results reported in this paper are available at https://github.com/bjbouman/cell-state-velocities.

## 4 Discussion

In this manuscript, we study some of the current challenges in the inference of cell state velocities from mRNA data and suggest novel approaches for tackling these problems. We argue that one of the interests in obtaining cell state velocities is to conserve information about variance between individual cells. This variance can inform us about fluctuations of the dynamics, cell plasticities and heterogeneity. We demonstrated that the processing procedure, several data smoothing steps and the visualisation approach in existing methods kill such biologically meaningful variance. The resulting information is closer to knowledge we could get from pseudotemporal ordering of cells than true velocity directions. The result is good looking cell velocity maps (i.e. conforming the expected pseudotime directions) that do not reflect the reality of the information contained in the u-s mRNA data.

We propose and design the *κ*-velo approach, which addresses the relative scaling of velocity components across genes combined with consistent processing and visualisation approaches. *κ*-velo’s velocity components’ scaling is based on the assumption that cell densities can be used as a proxy of typical travel time between two cell states. Although heterogeneous cell birth and death rates along the differentiation path disturb this assumption, we demonstrate how our model achieves better estimation of velocities than current methods on the pancreas data, showing a more plausible range of splicing rates and velocity magnitudes in several differentiation regions. To improve this model, one could consider estimating the heterogeneous cell birth and death rates based on the activity of apoptotic and proliferation genes [21]. This could account for how this would affect cell densities and thus the inferred travel times in future work. Alternative proxies of time measurement, such as diffusion distance, could also be designed instead of our cell density based approach. In Section 3.3, we found that there can be difficulty in fitting reaction rate parameters for genes that do not display evident kinetics on the u-s phase trajectory. Here, the method could be improved by including a prior on the expected kinetics during fitting. For example, prior knowledge about the pseudotemporal order of clusters could be included during the assignment of cells to transcriptional states of induction or repression.

For applications in which a general overview of a cell population’s directed motion in the expression state suffices, we propose the eco-velo heuristic approach. This approach eliminates multiple cumbersome steps of the standard methods, such as strict gene set selection, imputation and parameter fitting, all of which are computationally expensive and can kill some of the signal.

We also raise awareness about the time scales for which average velocities are being estimated. It would be interesting to measure velocities at multiple time scales for getting an overview on the “plans” individual cells have in preparation for their short- or long-term developmental journey. This supports growing interest for inferring cell state velocities from other pairs of single-cell data modalities, e.g. gene expression level coupled with chromatin accessibility or mRNA coupled with protein levels that correspond to different time scales of gene regulation. Furthermore, inferring cell state velocities from modalities in which measurements are more accurate (in comparison to the uncertainty in quantification of unspliced-spliced mRNA counts) can enhance our ability to understand the biological variation in cell state velocities rather than variations due to measurement noise.

To conclude, we suggest that a comprehensive grasp of what we are actually estimating and visualising as cell state velocities is crucial for obtaining a full description of cell differentiation dynamics. True cell state velocities encompass both stochastic and deterministic parts of the biological dynamics. This information can be complementary to attempts for describing cell differentiation as a full diffusion process [1, 5, 9, 22, 23, 24, 25] which contains the three terms of deterministic, stochastic and cell birth and death rates. Reliable quantification of cell state velocities in different transcriptional regions can put the relative magnitude (i.e. coefficients) of these terms into perspective in relation with one another.

## Acknowledgement

We would like to thank Max Delbrück Center for Molecular Medicine as well as the Bundesministerium für Bildung und Forschung (BMBF) (grant for ‘junior consortia in systems medicine’) for funding this study.

