## Supplementary notes and figures for "Towards reliable quantification of cell state velocities"

**2 Supplementary Figures** **6**

**References** **16**

### 1 Supplementary notes

#### Note 1: Kappa from s(t)

Let $\Delta t_{ij}$ be a measure of time that can be used to relate time between two states $i, j$ across genes, with $i$ before $j$ in time. Consider one gene with true parameters of reaction rate $\theta = (\kappa\alpha, \kappa\beta, \kappa\gamma)$ and recovered parameters $\theta = (\alpha, \beta, \gamma)$.
The equation for spliced (s) counts as a function of time is $s(t) = s_0 \exp(-\kappa\gamma(t - t_0)) + \frac{\alpha}{\gamma}(1 - \exp(-\kappa\gamma(t - t_0))) + \frac{\alpha - \beta u_0}{\gamma - \beta}(\exp(-\kappa\gamma(t - t_0)) - \exp(-\kappa\beta(t - t_0)))$, where $s_0, u_0$ are the initial condition of the s and unspliced (u) counts at time $t_0$. This yields for two measurements from cell i and j:

$$s_j = s_i \exp(-\kappa\gamma\Delta t_{ij}) + \frac{\alpha}{\gamma}(1 - \exp(-\kappa\gamma\Delta t_{ij})) + \frac{\alpha - \beta u_i}{\gamma - \beta}(\exp(-\kappa\gamma\Delta t_{ij}) - \exp(-\kappa\beta\Delta t_{ij})) \quad (1)$$

Solving for $\kappa\Delta t_{ij}$ we get:

$$\kappa\Delta t_{ij} = -\frac{1}{\gamma} \log \frac{s_j - \alpha/\beta + \frac{\alpha - \beta u_j}{\gamma - \beta}}{s_i - \alpha/\beta + \frac{\alpha - \beta u_i}{\gamma - \beta}} \quad (2)$$

### Note 2: Recovery of kappa from cell densities

For one gene fitted by scVelo [1], we recover the parameters of reaction rate $\theta = (\alpha, \beta, \gamma)$, $t(i)$ the time assignment for cell $i$, and $u_{t(i)}$ the assigned unspliced counts for $i$.
Given two cells $i, j$ that are in the same state $k$ of transcriptional induction or repression, and $i$ happens before $j$. We count $d(i, j) = n$ the number of cells in state $k$ with assigned counts between $u_{t(i)}$ and $u_{t(j)}$. The equation for $\kappa$ gives us:

$$\kappa n = -\frac{1}{\beta} \log \frac{u_{t(i)} - \alpha/\beta}{u_{t(j)} - \alpha/\beta} \tag{3}$$

$$\kappa d(i, j) = f(i, j) \tag{4}$$

Without measurement noise this equation should return the same value for any pairs of cells in the same transient state of transcription. To account for noise, we randomly sample pairs of cells $i, j$ that are in the same transcriptional state, and compute $d(i, j)$ and $f(i, j)$ for each pair. Plotting $d$ on the $x$-axis and $f$ on the $y$-axis, the slope of the corresponding line gives us $\kappa$ (see Figure S2B). Towards the extremes if one (or both) of $i, j$ is in steady-state and falsely assigned to a transient transcriptional state, $f$ will be smaller than expected from equation 4, and we will have points right of the $\kappa$ line on the $d$, $f$ plot (see Figure S2B). The $\kappa$ line is still given by the left slope of the $d$, $f$ plot. To recover $\kappa$, we fit a parallelogram to the points such that the area of the parallelogram is minimal while maximising the number of points in the parallelogram (see Figure S2B), the $\kappa$-slope is then given by the left side of the parallelogram.
Note that the recovered $\kappa$ for all genes are still off by a same constant factor $c$, that would relate the density to true time. Since this factor is constant for all $\kappa$, we can still relate velocities across genes.

### Note 3: Projection of velocities onto PCA embedding

For projection of end of the velocity arrows (test set data points) onto the existing principal component analysis (PCA) of initial cell positions (training set), we use the weight matrix $W_{g\times k}$ that transformed the training points $S_{n\times G}$ into the PCA transformation $Y_{n\times k}$ of $S$ on the first $k$ principal components. We have:

$$Y = SW \tag{5}$$

Since PCA is a linear transformation of gene space, we can apply the same transformation on the future cell states $S + \vec{V}$. The PCA-transformed future states are:

$$Y_{fut} = (S + \vec{V})W \tag{6}$$

The single-cell velocities can then be visualised on the first $k$ principal components.

### Note 4: Projection of velocities onto nonlinear embedding

In nonlinear projection methods, s.a. diffusion maps [2], t-SNE [3] and UMAP [4], a transition probability matrix $P_{n1\times n1}$ from the cell states $S_{n1\times G}$ is transformed into a low-dimensional embedding $Y_{n1\times k}$.
From the training set $S_{n_1\times G}$ , we get the row-normalised transition probability matrix $P_{n_1\times n_1}$ between all pairs of cells $i, j$:

$$P_{i,j} = \frac{1}{\tilde{Z}(i)} \frac{\exp(-\frac{-||S_i - S_j||^2}{2\sigma^2})}{Z(i)Z(j)} \tag{7}$$

With normalising factors:

$$Z(i) = \sum_{j \in \Omega} \exp(-\frac{-||S_i - S_j||^2}{2\sigma^2})$$

$$\tilde{Z}(i) = \sum_{j \in \Omega} \frac{\exp(-\frac{-||S_i - S_j||^2}{2\sigma^2})}{Z(i)Z(j)}$$

The Gaussian width $\sigma^2$ determines the length scale over which each cell can randomly diffuse. Note that in this matrix the diagonal will be the maximal value of each row, and not set to zero $P_{i,i} \neq 0$. We can recover a transformation $W_{n1 \times k}$ from $P$ to $Y$ given by:

$$W = P^{-1}Y \tag{8}$$

To project the future states onto the existing embedding, we first compute the transition probability matrix $P'_{n2 \times n1}$ from the future states to the old ones, and apply the transformation $W$ to $P'$:

$$Y_{fut} = P'W \tag{9}$$

Note that the original transformation from the training set is not linear. $W$ is an approximation of the original transformation that will only yield meaningful results if the newly introduced points $S + \vec{V}$ are within the original training manifold $S$. This should hold if $\vec{V}$ is small.

#### Diffusion maps

In diffusion maps the embedding $Y_{n1 \times k}$ corresponds to the first $k$ right eigenvectors $Y = [\psi_0, \ldots, \psi_k]$ of $P$. With the corresponding ordered eigenvalues $\boldsymbol{\lambda} = \lambda_0 \geq ... \geq \lambda_k$, we have: $PY = \lambda Y$. Diffusion maps are a special case of eq. 8 where the eigenvector matrix $Y$ and eigenvalues $\lambda$ give us the transformation:

$$Y_{fut} = (P'Y)/\lambda \tag{10}$$

### Note 5: Processing of datasets

The RNA velocity workflow truly starts with counting the spliced and unspliced reads (preprocessing), followed by several operations (filtering, normalisation, etc.) to prepare the data for the downstream analysis (processing). Both preprocessing and processing have major effects on the outcome of the downstream analysis. Therefore it is essential to understand the implications of each step in the (pre)processing pipeline. After careful investigation of the scVelo processing pipeline [1], we have come to the conclusion that some processing steps should be updated to reduce the introduction of artefacts and to improve the recovery of the rate parameters $\alpha$, $\beta$ and $\gamma$. Below, we will describe our proposed four step processing pipeline, which is part of the $\kappa$-velo workflow, and give a detailed explanation of the implications of each step. Additionally, we have three comments on processing, which are not addressed in the six steps, but which are nevertheless important to take into consideration.

#### Step 1: Filter HVGs

To decrease the dimensions of the single-cell dataset we select only those genes that have a high variability. We use analytic Pearson residuals to identify the variability of each gene [5]. We prefer this method over other methods, because it recovers both low- and high-expression genes with high variance. The HVGs are identified using the spliced counts only.

#### Step 2: Select genes with sufficient counts

After selecting the HVGs, we filter the genes once more to include only genes that have sufficient spliced and unspliced counts. In step 4 the spliced and unspliced counts for each gene in each cell are imputed using the nearest neighbours to decrease the noise in the measurements. Imputation can create artificial clouds of points in the unspliced/spliced portrait for some genes with extremely low unspliced and/or spliced counts, for which the dynamics cannot be reliably recovered (see Figure S9). Therefore, we filter out all genes for which the spliced and/or unspliced counts are below a certain threshold. For the dataset of pancreatic endocrinogenesis, all genes with a maximum spliced or unspliced count below 4 were filtered out for further downstream analyses.

#### Step 3: Normalisation

In the processing pipeline of scVelo, unspliced and spliced counts are normalised separately. Normalisation is used to reduce the effect of count depth differences between cells [6]. Most commonly, the counts in each cell are divided by the total counts for that cell and multiplied by a scaling factor. Since spliced and unspliced counts derive from the same cells, the total counts of each cell can be calculated by summing both for each cell. Rather than normalising using separate total counts, the spliced and unspliced counts should be normalised by their combined total counts. This proves especially important in datasets where cell types have different ratios of total spliced to total unspliced counts, such as the erythroid lineage in the mouse gastrulation dataset published by Pijuan-Sala et al. [7] (Figure **??**). In this erythroid branch Barile et al. found several genes with multiple rate kinetics (MURK genes) [8]. Those genes distinguish themselves in the unspliced/spliced phase portrait with a rapid increase of unspliced counts in the more differentiated cell types. Perform a combined normalisation in the original protocol, the curvature of these genes changes in the u-s phase portrait (Figure **??**), indicating that this could be an artefact of normalisation.

#### Step 4: Imputation

The imputation step in the pipeline averages the spliced and unspliced counts of each cell using its nearest neighbours. Since we filtered out a large proportion of genes in step 1 and step 2, the PCA used for calculating the nearest neighbours is recalculated. For the computation of the PCA coordinates, only the spliced counts are used. For PCA calculation, the spliced counts are scaled to stabilise the variances of the genes.

#### Removal of low-likelihood genes

Although a part of the velocity calculations, the $\kappa$-velo workflow includes on more filtering step. After running `scvelo.tl.recover_dynamics`, each gene is assigned a likelihood, which indicates how well a gene is described by the recovered phase trajectory. The genes can then be ranked according to likelihood of the fit. Only genes with a likelihood above a certain threshold are used for the downstream velocity analysis and embedding. Like step 2, the used threshold depends on the dataset.

#### Comment 1: False assignment of unspliced to spliced

The RNA velocity protocol is based on separating spliced from unspliced molecules. There exists multiple counting software to quantify these molecules. Here, we will focus on the quantification rules as described in [9]:

- Molecules are counted as "spliced" if all of the reads in the set supporting a given molecule map only to exonic regions.
- Molecule are counted as "unspliced" if all, or at least one read in the set spans an exon-intron boundary, or is mapped to an intron for every compatible transcript models.
- Molecules are counted as "ambiguous" if some of the compatible transcript models have exonic mappings, and others intronic mappings. These counts are disregarded in further analyses.

Since unspliced molecules also contain exonic regions and we do not measure the whole molecule we expect that some of the unspliced molecules will be misassigned to spliced. The recovered counts would then follow $(u_r, s_r) = (u_t - \eta u_t, s_t + \eta u_t)$, with $\eta$ the rate of misassignement and $(u_t, s_t)$ the unknown true unspliced and spliced counts for one cell. This will affect both the steady-state ratio, and the curvature of the gene-specific dynamics. Because of this, it comes as no surprise that the choice of counting software and parameters can affect downstream analysis [10].

#### Comment 2: Log transformation

Long-read sequencing methods such as smart-seq yield a better disentanglement of u and s counts, hence seem more suitable for RNA-velocity analysis. Whereas several current single-cell methods (PCA, t-SNE, nearest neighbours search, etc) are usually used for non-UMI data only after log-transformation, the u-s dynamical equations used for parameter estimation in the RNA-velocity framework are defined for raw counts (e.g. by using $u_0 = 0$ and $s_0 = 0$ as the initial point in the u-s phase portraits). Most likely, this is also the reason that scVelo does not log transform the spliced and unspliced counts that are used for recovery of the dynamics. Both of scVelo's functions `scvelo.pp.filter_and_normalize` and `scvelo.pp.log1p` do not log-transform the u, s layers used for parameter recovery. To avoid any inconsistency, our processing pipeline does not include any log transformation.

#### Comment 3: Batch correction

With the technological advance of single-cell technologies, more and more single-cell RNA seq datasets may be compiled from multiple experiments. In most multi-sample cases, a batch correction is essential to recover the biological variation, while removing batch-effects [11]. However, batches may affect unspliced and spliced counts differently. Therefore it is important the be cautious when using any batch correction, and to always check that the recovered dynamics do not contain batch- or batch correction-related artefacts.

### Note 6: Simulation

For the simulation we randomly sampled $G$ log-normally distributed parameters $\theta = (\alpha, \beta, \gamma)$ with $\log(\theta) \sim N(\mu, \Sigma)$, $\mu = (1, 0.2, 0.05)$ and $\Sigma_{ii} = 0.04, \Sigma_{ij,i \neq j} = 0.04 * 0.02$. The parameters are scaled by a scaling factor $\kappa$: $\theta = (\kappa\alpha, \kappa\beta, \kappa\gamma)$. The $n$ cells follow a uniformly distributed hidden true time $t$, with cell-specific time points $(t_1, ..., t_n)$. The unspliced and spliced counts for genes are simulated following

$$u(t) = u_0 \exp(-\beta\tau) + \frac{\alpha}{\beta}(1 - \exp(-\beta\tau))$$
$$s(t) = s_0 \exp(-\gamma\tau) + \frac{\alpha}{\gamma}(1 - \exp(-\gamma\tau)) + \frac{\alpha - \beta u_0}{\gamma - \beta}(\exp(-\gamma\tau) - \exp(-\beta\tau))$$

with $\tau = t - t_0 - t_s$, $t_0$ the time of transcriptional switch (on to off) and $t_s$ the time of transcriptional start (off to on). The time point of transcriptional start for a gene was chosen inversely to the speed of the gene so that all genes reach the end of transcription at the same time. This results in a trajectory with high plasticity at the beginning when fast genes are not yet activated and lower plasticity at the end when all genes are in transient state.
Random normal noise was added to the unspliced and spliced counts.

### Note 7: Evaluation of inferred velocities in simulation data

We use two measures to evaluate the difference between two vectors $\vec{v}_t$ and $\vec{v}$ in PCA space. The cosine similarity $\frac{\vec{v}_t \times \vec{v}}{\|\vec{v}_t\| \|\vec{v}\|}$ is the cosine of the angle between the two vectors. The difference in vector length is given by as $\|\vec{v}_t\| - \|\vec{v}\|$.

### 2 Supplementary Figures

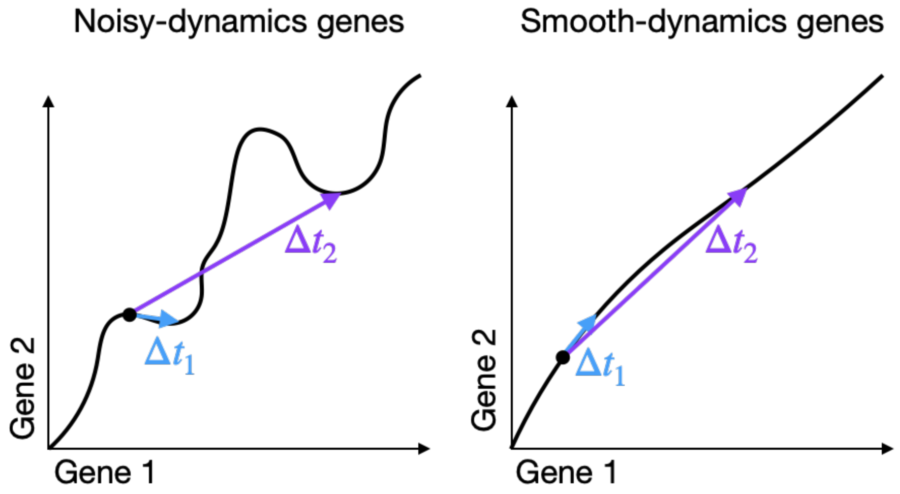

**Figure S1:** Average velocities for different time scales can be very different if the expression dynamics are not smooth. On the left is the example of two noisy genes: the average velocity over $\Delta t_1$ is very different from the average velocity over $\Delta t_2$. For smooth gene dynamics as shown on the right, the average velocities are more similar.

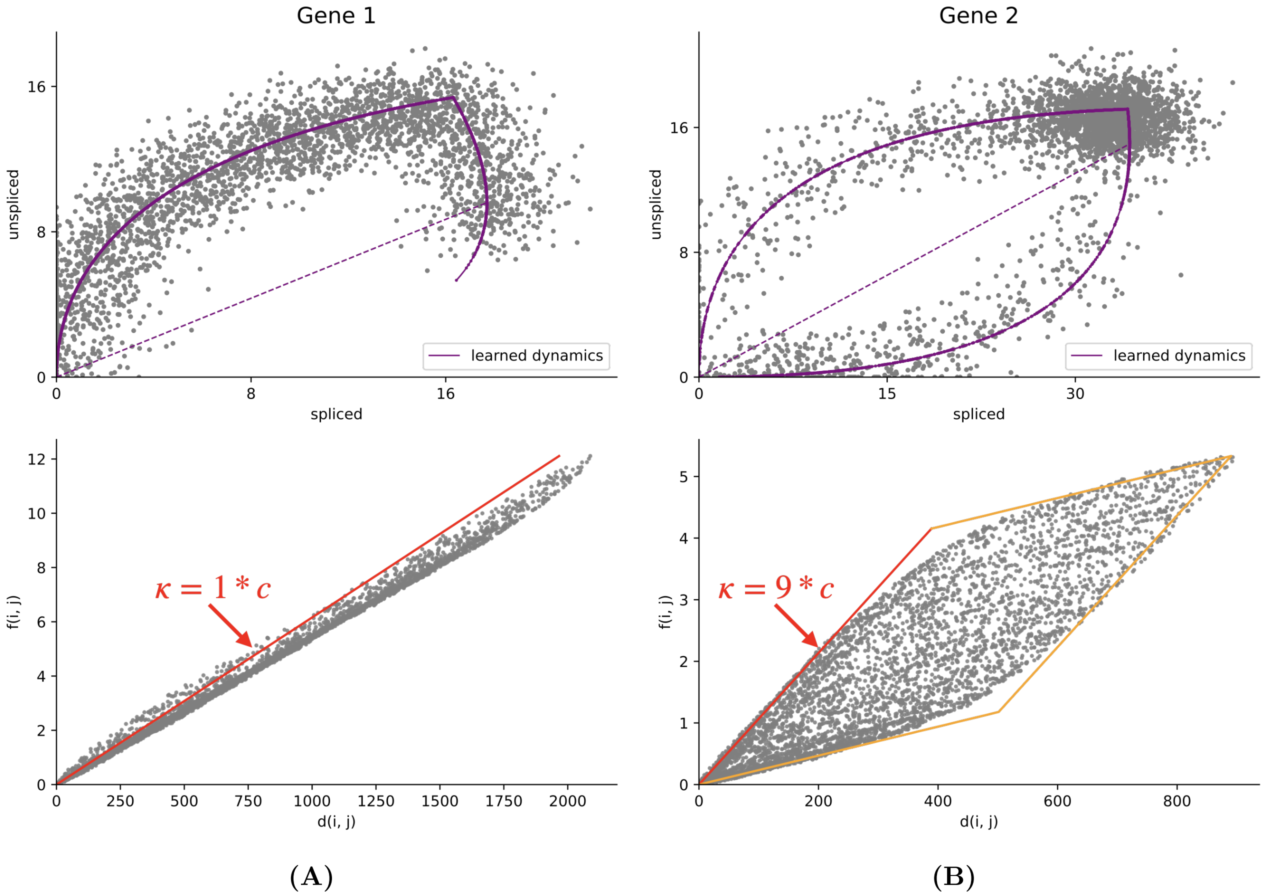

**Figure S2:** Density estimation for two simulated genes with different time scales. $c = 10^{-3}$ is a constant scaling factor. The two simulated genes have the same reaction parameters $\theta$ but those for gene 2 are scaled by 10. (A) a slow gene, where no cells are in steady-state. The slope of the line gives us $\kappa_g 1$ directly. (B) a fast gene, where a lot of cells are in steady-state. The slope of the red line gives us $\kappa_{g2}$.

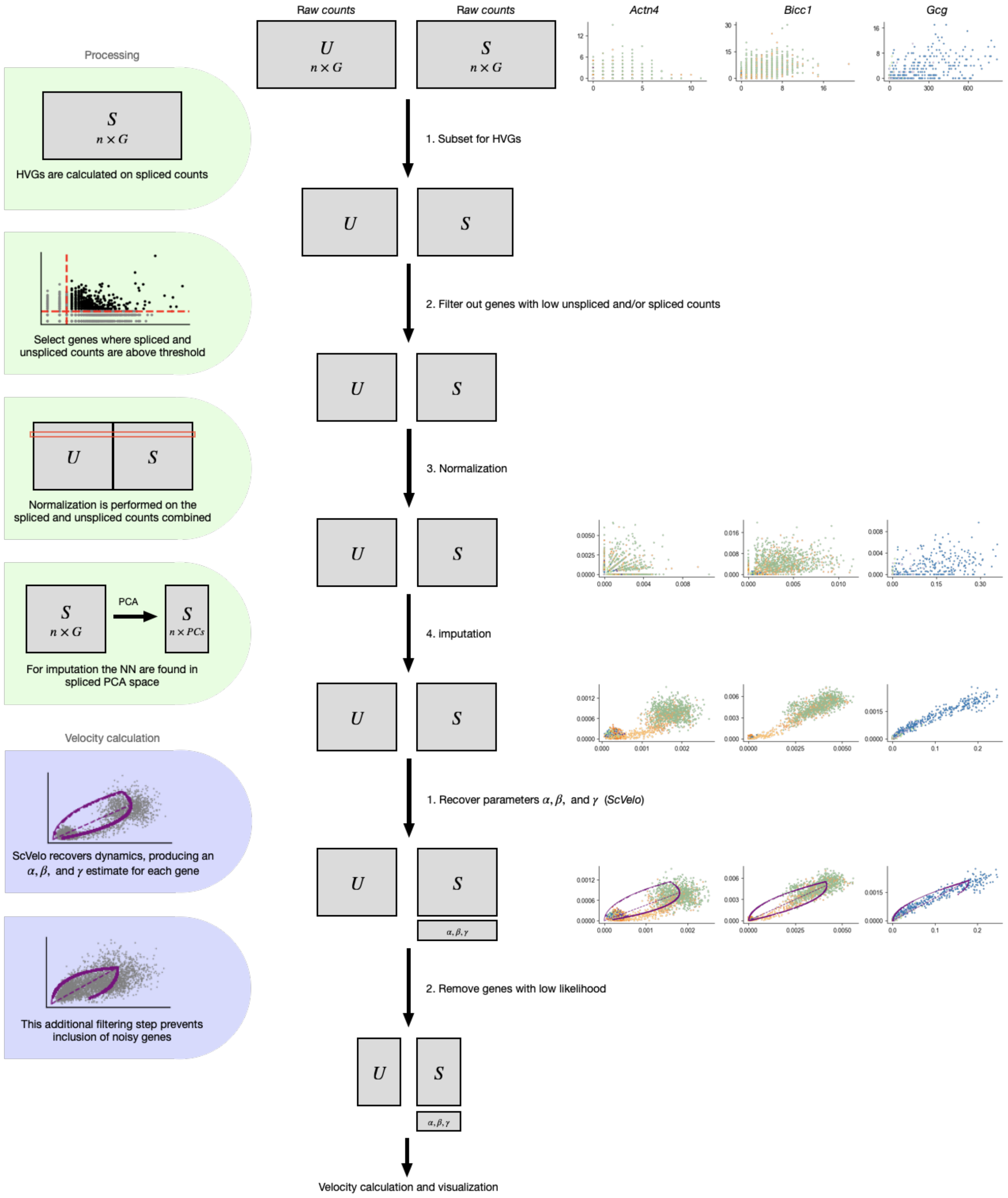

**Figure S3:** Workflow of processing and part of velocity calculation. In the middle, a schematic representation of how the spliced and unspliced matrices change during each step is shown. On the right, the u-s phase portraits of genes *Actn4*, *Bicc1* and *Gcg* (from the pancreas endocrinogenesis dataset) to illustrate how the spliced and unspliced counts change after each step.

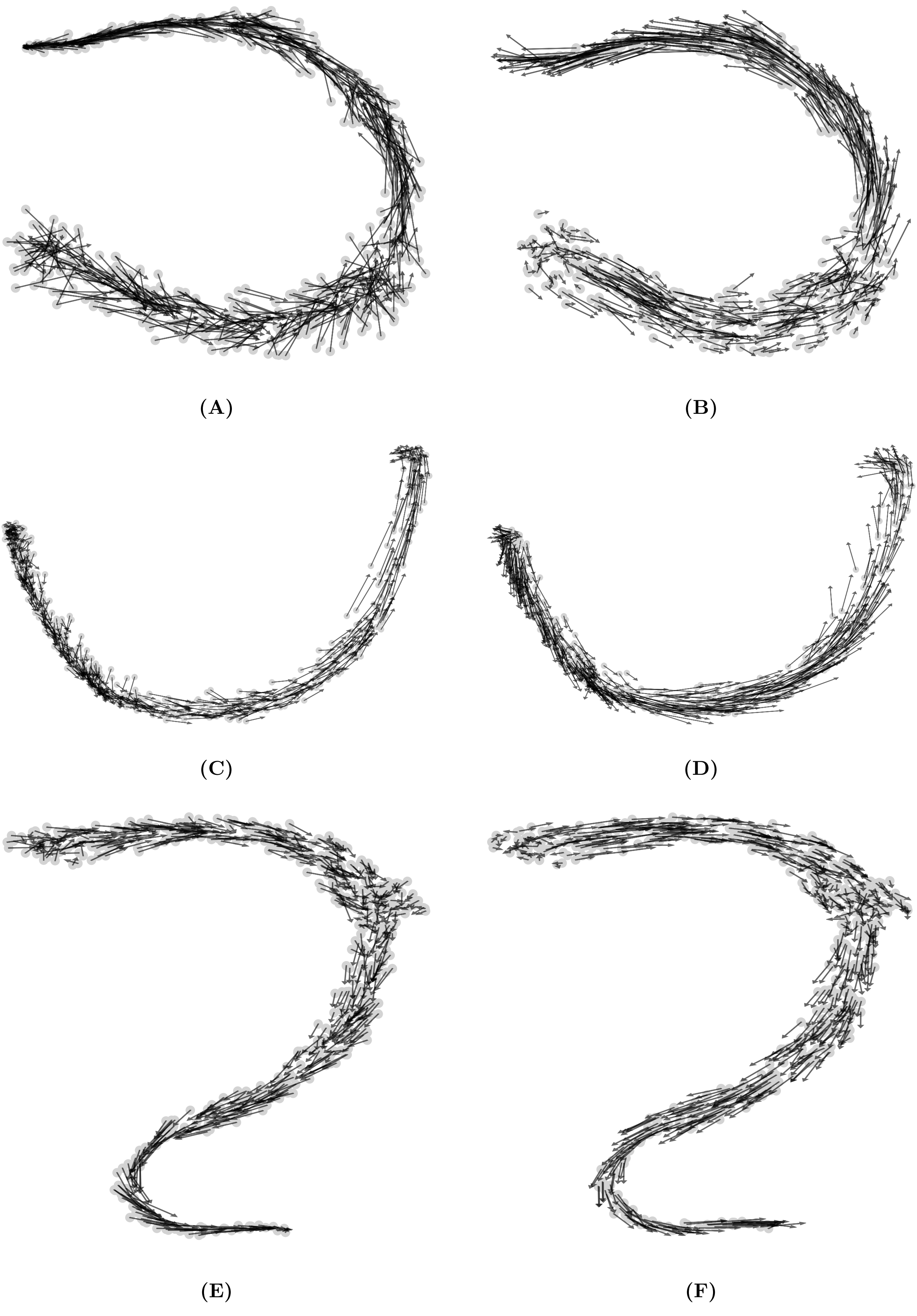

Figure S4: Projection of the velocity arrows (test set data points) onto existing embedding of initial cell positions (training set). We compare our projection approach (left column) to scVelo's [1] (right column) embedding for t-SNE [3] in (A) and (B), diffusion maps [2] in (C) and (D), and UMAP [4] in (E) and (F).

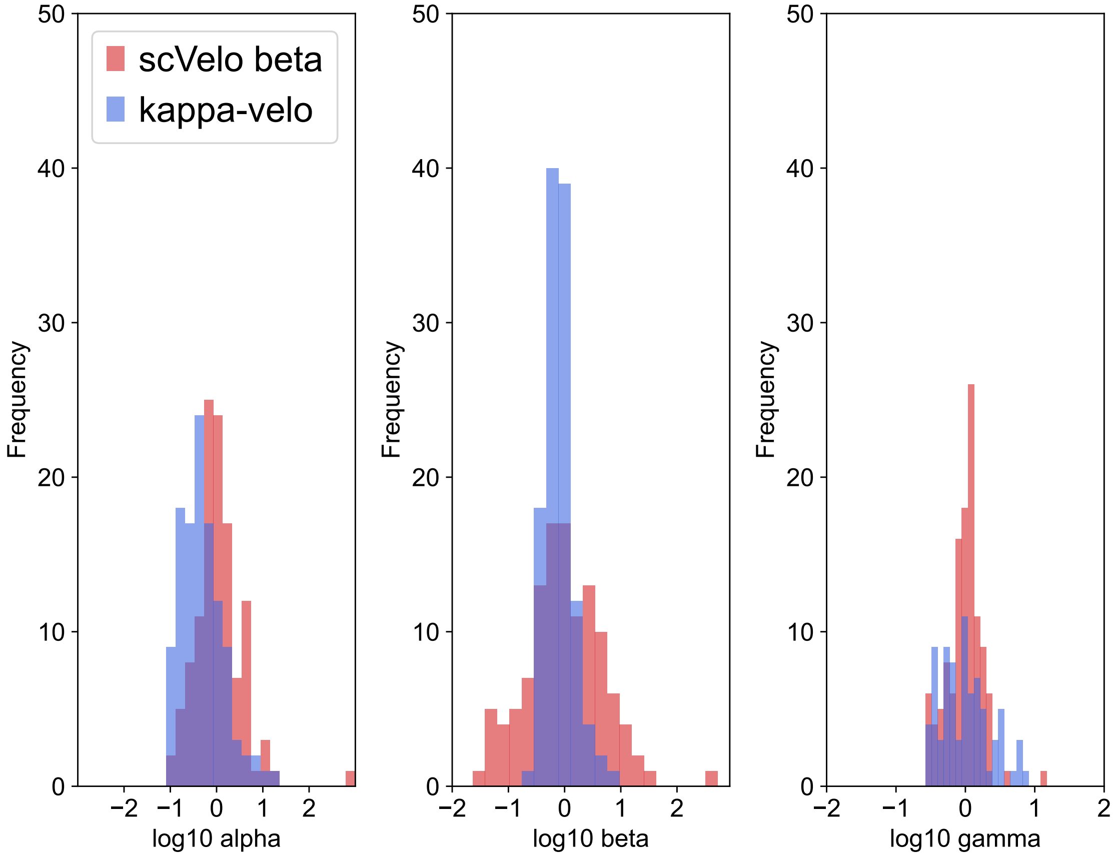

**Figure S5:** Range of transcription rate $\alpha$, splicing rate $\beta$, and degradation rate $\gamma$ estimated by scVelo (in red) and $\kappa$-velo (in blue).

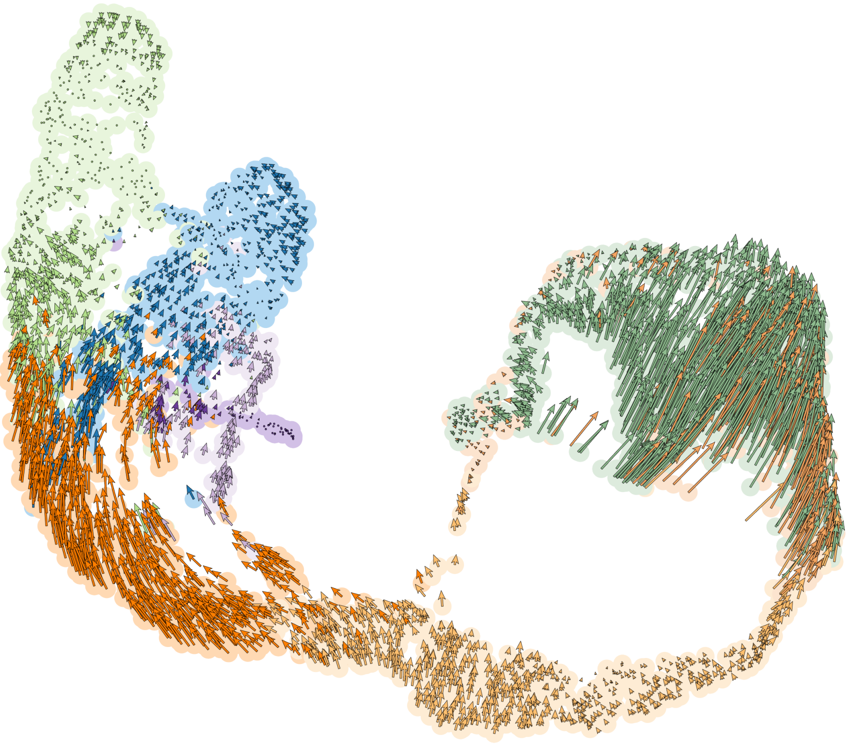

**(A)**

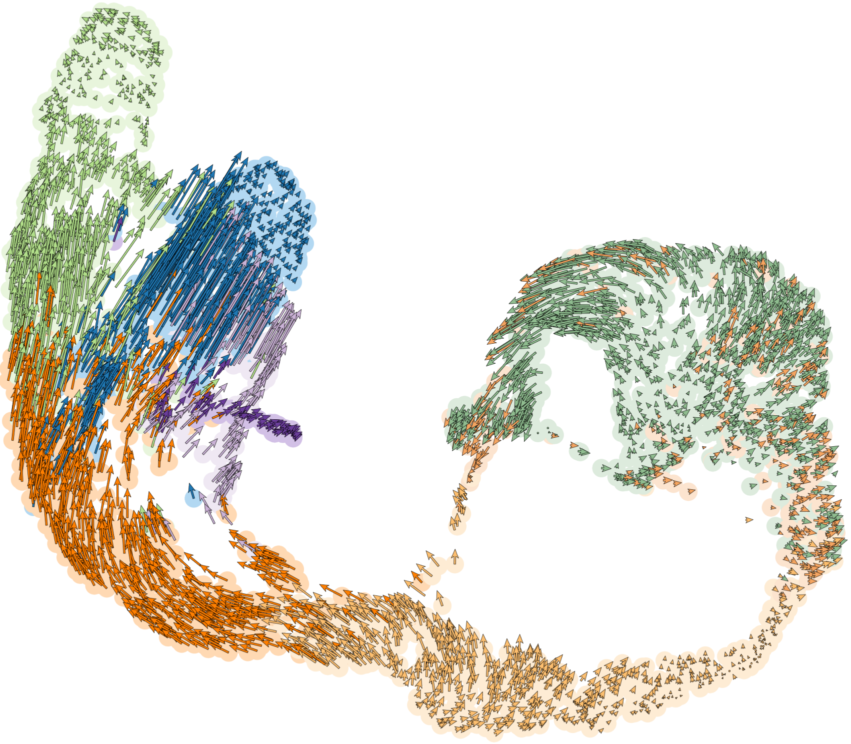

**(B)**

**Figure S6:** Smoothed $\kappa$-velo projection of velocities in the pancreas endocrinogenesis dataset. The two UMAPs compare scaled (A) and non-scaled (B) velocities. Velocities were smoothed by averaging over the 50 nearest neighbours. Neighbourhoods are calculated in S space.

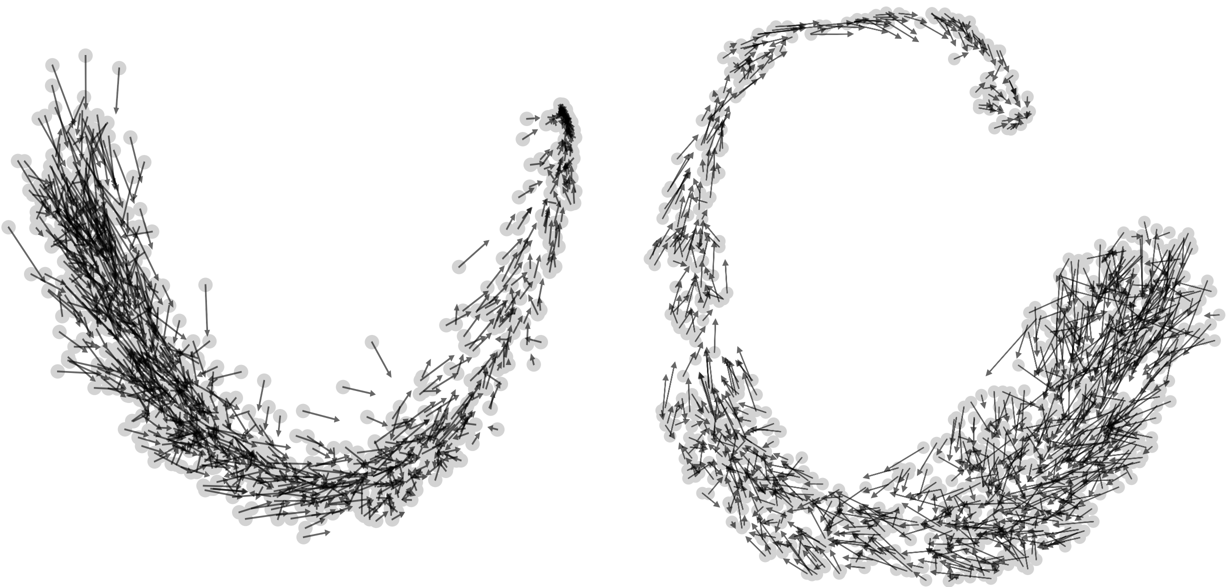

**(A)**

**(B)**

**Figure S7:** eco-velo projection of velocities (calculated on simulations) shown on PCA in (A) and UMAP [4] in (B).

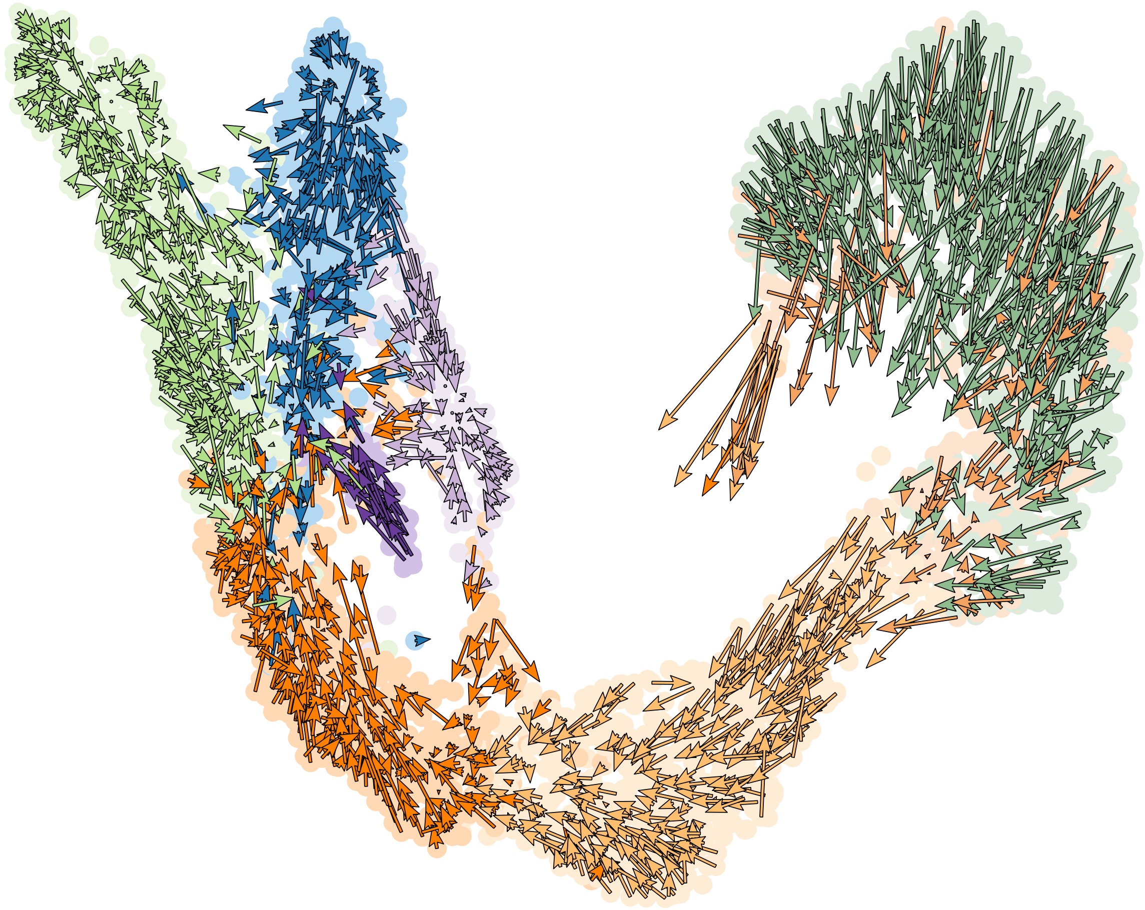

**Figure S8:** Smoothed eco-velo projection of velocities in the pancreas endocrinogenesis dataset. Velocities were smoothed by averaging over the 50 nearest neighbours. Neighbourhoods are calculated in S space.

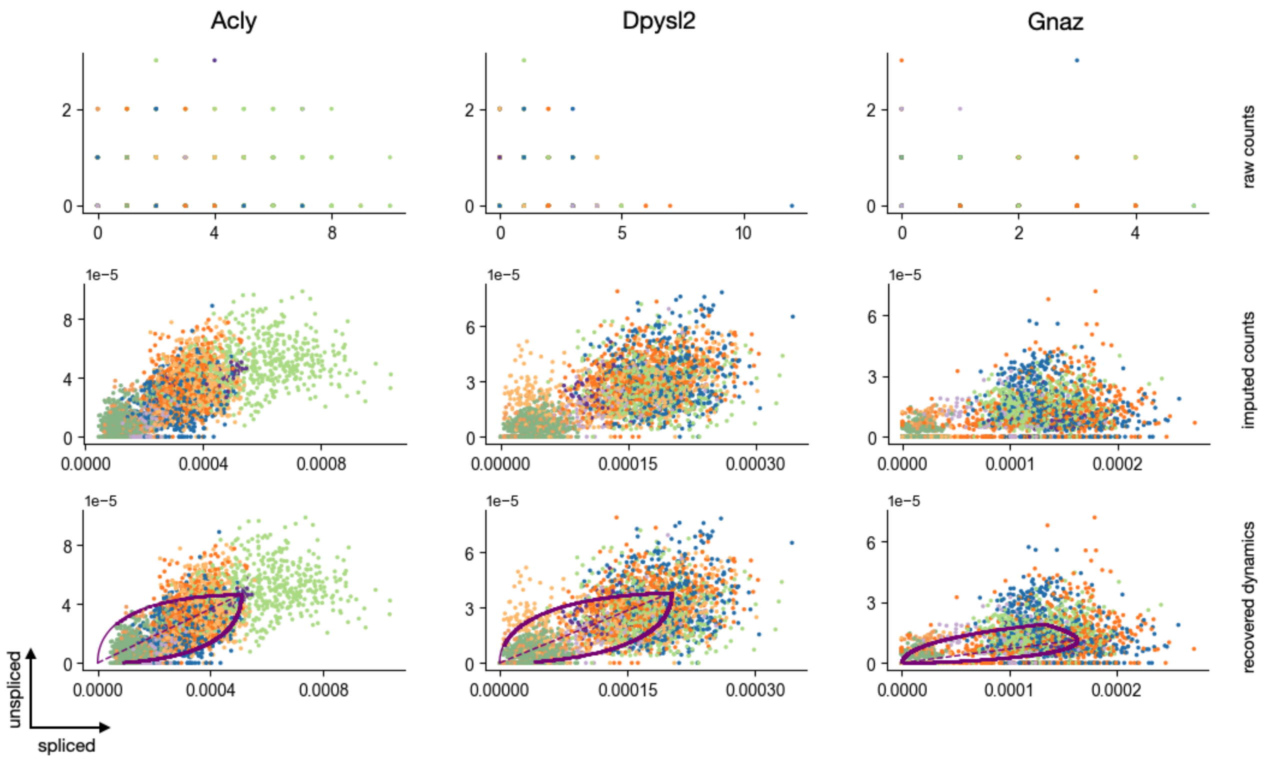

**Figure S9:** The u-s phase portrait of *Acly*, *Dpysl2* and *Gnaz* (from the pancreas endocrinogenesis dataset), which are all genes with insufficient unspliced counts. Here, we show how scVelo would recover the dynamics if these genes were not filtered out.

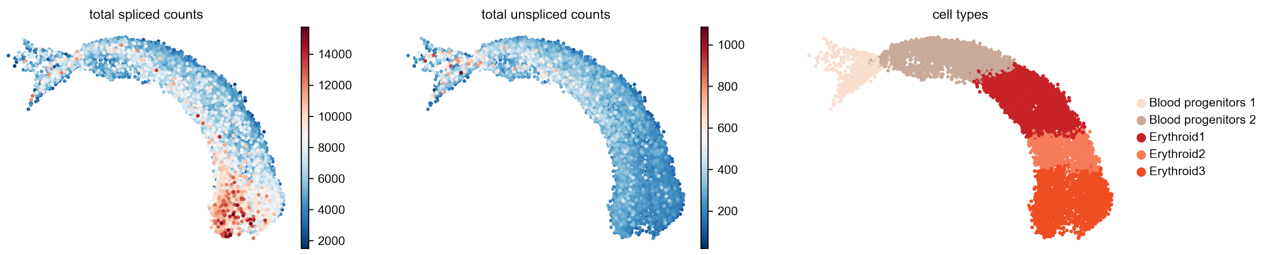

**(A)**

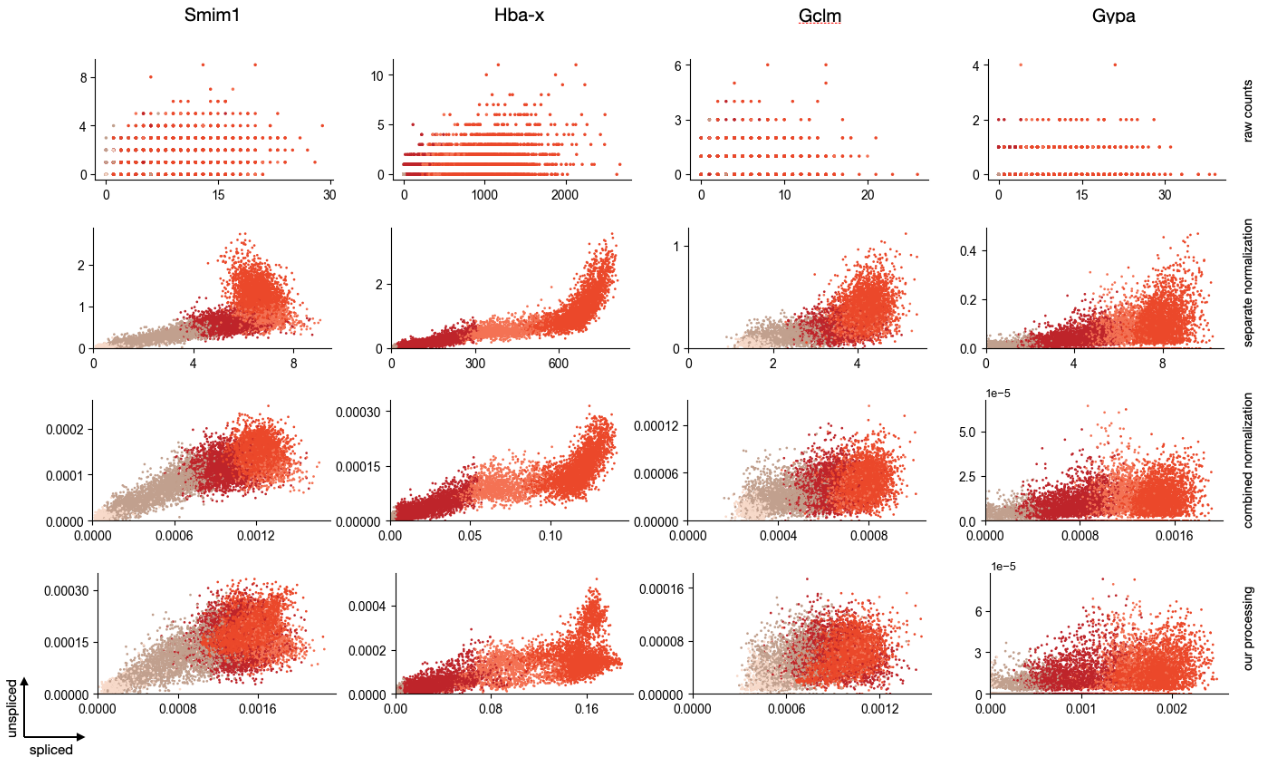

**(B)**

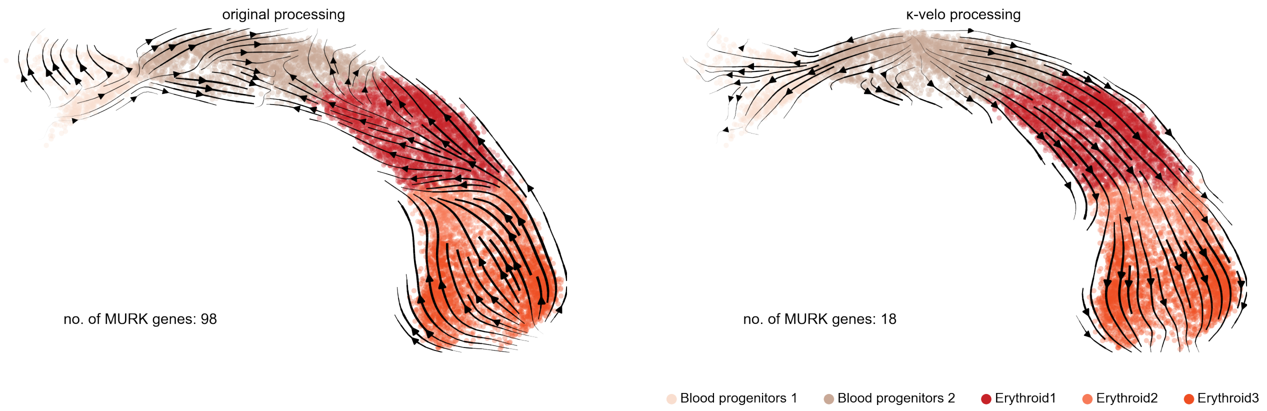

**(C)**

**Figure S10:** The scRNA-seq dataset on the erythroid lineage of mouse gastrulation [7] has been described in the context of RNA velocity by Barile et al. [8]. Here, we show that the subset has a varying ratio of total unspliced to total spliced counts in different cell types (A). This results in artefacts when using the standard scVelo processing pipeline (U and S normalised separately) (B, second row). Those artefacts are mostly resolved by normalising U and S combined (B, third row), which is part of the $\kappa$-velo processing workflow (B, last row). Using the $\kappa$-velo processing workflow fixes some of the reported de-differentiation (C).

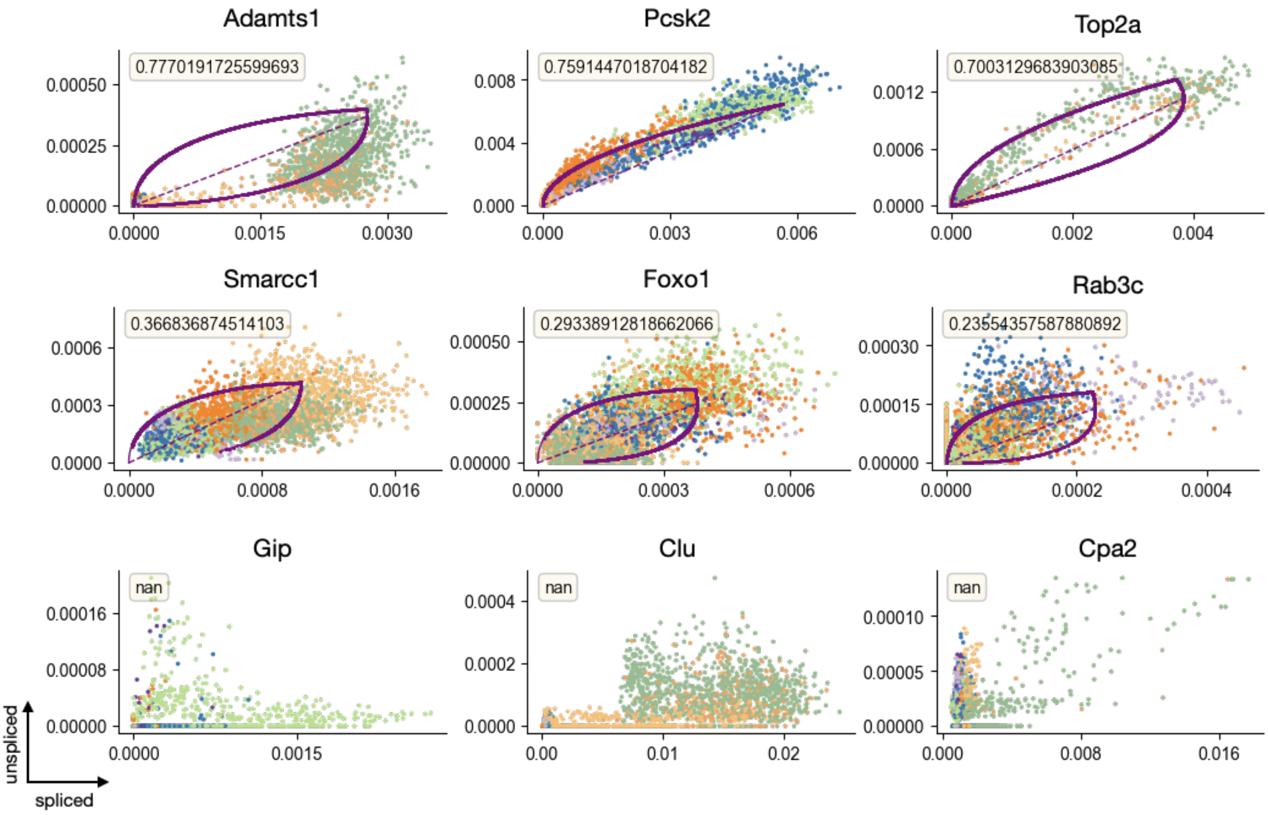

**Figure S11:** Recovered dynamics (purple) and likelihood (indicating how well the recovered dynamics fit the measured data points) for nine different genes from the pancreas endocrinogenesis dataset. On the top row three genes with high likelihood, which will be considered for the downstream velocity calculation are shown. On the second row, three genes with a low likelihood (below 0.4), which will be removed are shown. On the bottom row, three genes for which scVelo did not recover any dynamics and therefore will also be removed for further downstream purposes are shown.
